# Galacto-oligosaccharides modulate the juvenile gut microbiome and innate immunity to improve broiler chicken performance

**DOI:** 10.1101/631259

**Authors:** Philip J. Richards, Geraldine M. Flaujac Lafontaine, Phillippa L. Connerton, Lu Liang, Karishma Asiani, Neville M. Fish, Ian F. Connerton

## Abstract

Improvements in growth performance and health are key goals in broiler chicken production. Inclusion of prebiotic galacto-oligosaccharides in broiler feed enhanced the growth rate and feed conversion of chickens relative to a calorie-matched control diet. Comparison of the cecal microbiota identified key differences in abundance of *Lactobacillus* spp. Increased levels of *L. johnsonii* in GOS-fed juvenile birds at the expense of *L. crispatus* was linked to improved performance (growth rate and market weight). Investigation of the innate immune responses highlighted increases of ileal and cecal IL-17A gene expression counterposed to a decrease in IL-10 and IL-17F. Quantification of the autochthonous *Lactobacillus* ssp. revealed a correlation between bird performance and *L. johnsonii* abundance. Shifts in the cecal populations of key *Lactobacillus* spp. of juvenile birds primed intestinal innate immunity without harmful pathogen challenge.

**IMPORTANCE:** Improvements in the growth rate of broiler chickens can be achieved through dietary manipulation of the naturally occurring bacterial populations whilst mitigating the withdrawal of antibiotic growth promoters. Prebiotic galacto-oligosaccharides (GOS) are manufactured as a by-product of dairy cheese production, which can be incorporated in the diets of juvenile chickens to improve their health and performance. This study investigates the key mechanisms behind this progression and pin points *L. johnsonii* as a key species that facilitates the enhancements in growth rate and gut health. It also relates the role of the innate immune system in the response to the GOS diet.

## INTRODUCTION

The production of poultry for both meat and eggs has been increasing rapidly throughout the world (1) and the global poultry sector is expected to continue to grow as a result of growing population, rising income and urbanization (2). In this context, animal performance and feed conversion efficiency of fast-growing birds are decisive to the economic profitability of poultry meat production. Broiler chicken production is more sustainable and has a relatively lower environmental impact than other meat-based protein production (3). There have been massive increases in the growth rate and feed efficiency of broiler chickens since the 1940s, achieved largely through selective breeding and feed optimization (4). It is generally recognized that increases in performance are slowing as the advances made possible through these approaches are reaching their biological limit (5). The inclusion of antimicrobial growth promoters (AGPs), a practice banned in the European Union since 2006, is another way in which gains in productivity have been realized. The EU ban was imposed due to increasing concerns regarding the development of antimicrobial resistance and the transference of antibiotic resistance genes from animal to human microbiota (6). Although still widely used there have been reductions in the therapeutic use of antimicrobials in poultry production, which have led to an increase in intestinal health problems (7). To mitigate the effect of antimicrobial reduction a variety of strategies has been evaluated (8). These include the addition of dietary prebiotic (9, 10), phytobiotic dietary additives (11), the incorporation of beneficial enzymes in poultry feed (12), and the administration of live probiotic bacteria in various combinations of the above (7, 13). Recent developments in sequencing technologies have led to a greater understanding of mechanisms and effects of these treatments on the gut microbiota and the interaction with host-related functions involved in intestinal health (14, 15). It has been proposed that further improvements to broiler performance could be sought through deliberate cultivation of a beneficial gut microbiota in early development (7, 16). These bacteria are preferably autochthonous and mutualists in association with each other and their host.

Galacto-oligosaccharides (GOS) are non-digestible carbohydrates that have been shown to promote beneficial autochthonous bacteria, such as *Bifidobacteria*, *Bacteroides* and *Lactobacillus*. GOS are synthesized from lactose by *β*-galactosidase catalyzed transglycosylation to create molecules of differing length and linkage type (17, 18). Several studies have reported that GOS can improve the performance of poultry and produce profound differences in desirable bacterial groups inhabiting the gut (19, 20).

Maintenance and enhancement of gut health is essential for the welfare and productivity of animals (21). In addition to nutrient digestion and absorption, the intestinal mucosa constitutes a physical and immunological protective barrier for the integrity of the intestinal tract (22). Mutualistic commensals with immunomodulotary effects (autobionts) affect the development and function of various immune cell populations such as regulatory T-cells (Treg), Th17 T-helper cells, IgA-secreting plasma cells, natural killer cells (NK), macrophages, dentritic cells (DCs), innate lymphoid cells (ILCs) (23). Interleukin-17-producing CD4+ T lymphocytes (Th17 cells) contribute to host defense against pathogens and maturation of the immune response at an early age. Regulatory T-cells play critical roles in immune suppression (24, 25) thus optimum health is achieved through a balanced regulation of expression between Th17 cells and Tregs.

There is little systematic information regarding the interaction between prebiotic diet, performance, structure of gut microbiota and the host gene expression in poultry. In this study, the impact of a GOS diet was assessed in broiler chickens, by comparing a control diet fed cohort and a GOS diet fed cohort, from day of hatch until 35 days of age, corresponding to a typical commercial farm rearing period. Ancillary dietary trials were carried out to confirm the reproducibility of the beneficial effects. The innate immune responses to the two diets were assessed in ileal and cecal tissue biopsies by quantification of the relative expression of cytokine and chemokine gene transcription. Analysis of metagenomic profiles of GOS fed birds enabled the identification and isolation of autochthonous synbiotic organisms. Characterization of these isolates allowed an in-depth analysis of the effect of the GOS diet and synbiotic species abundance on bird performance and gut health.

## RESULTS

### Galacto-oligosaccharide (GOS) supplementation improves the growth performance of broiler chickens

Chickens fed GOS diet performed better than with those fed control diet, increasing in weight by 87.78g /day compared to 76.3g /day for control birds (*p* = 0.012; Figure 1A). The increased growth rate was apparent from an early stage in the rearing cycle (8 da) and correspondingly the slaughter age (day 35) weights of GOS fed birds in Trial 1 were significantly greater than that of control birds (*p* = 0.03; Figure 1B). Ancillary trials were carried out to demonstrate reproducibility with the birds housed in individual cages (Trial 2; Figure 1C) or group-housed (Trial 3; Figure 1D). The enhanced performance on the GOS diet was also evident in Trial 2 (*p* < 0.001) and Trial 3 (*p* = 0.057). Although Trial 3 marginally failed to meet significance, in this case the control diet birds performed exceptionally well with respect to live weight at day 35, but even under these circumstances the trend remained the same with the GOS diet producing beneficial effects. The feed conversion ratios (FCR) varied between trials but were reduced for birds on the GOS diets compared to the corresponding control groups (Figure 1).

**Figure 1.**
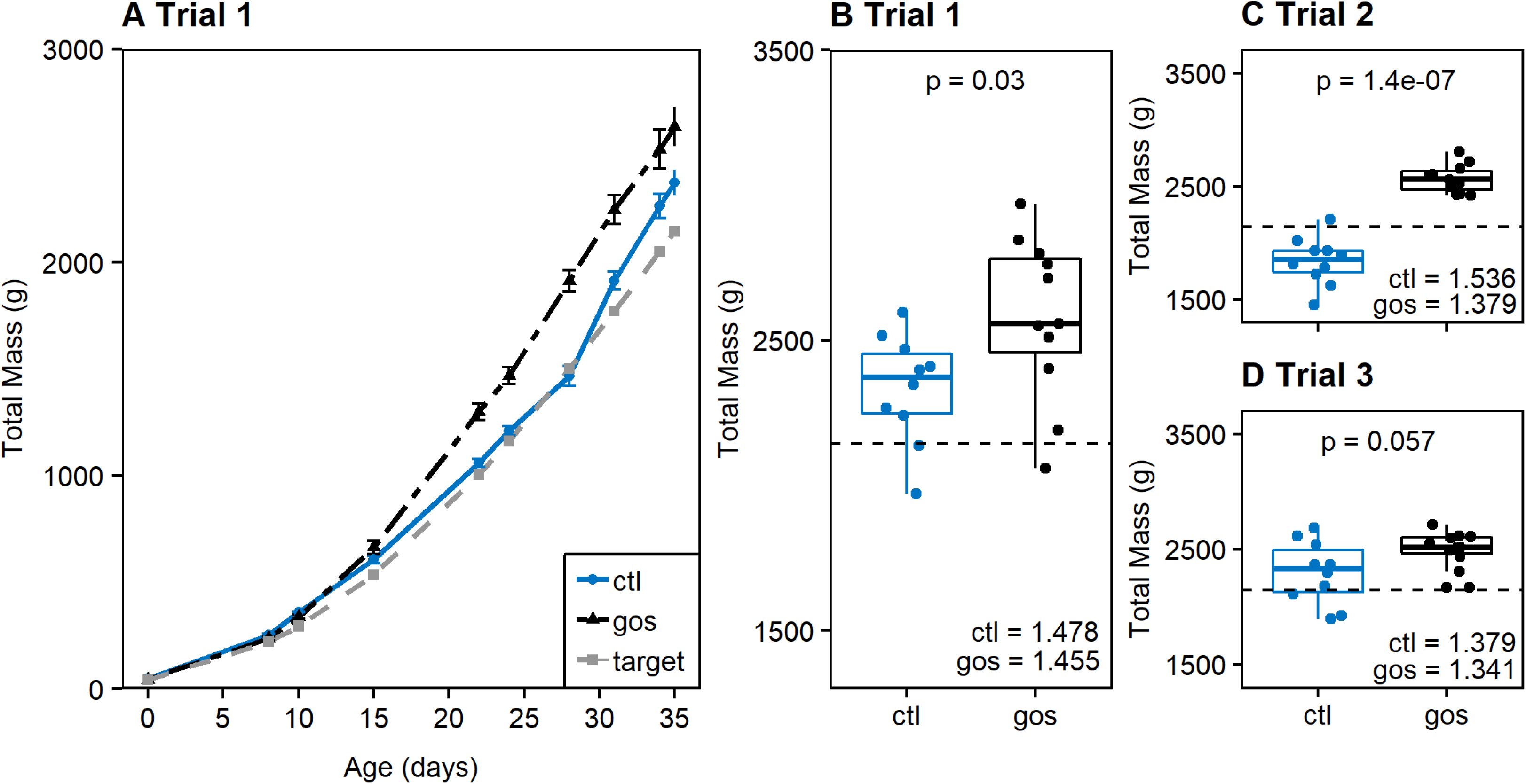
Galacto-oligosaccharide diet improves the growth performance of broiler chickens. (A) GOS diet trial comparing body weight of birds fed GOS diet (black) with birds fed control (ctl) diet (blue). The male Ross 308 broiler chicken performance objective weight progression (target; Aviagen Ross 308 Performance Objectives 2014) is indicted by the grey dashed line. (B) Box whisker plots showing the spread of data presented for 35 da trial 1. The discontinuous horizontal black line indicates the Ross 308 performance objective at 35 da for reference in each panel. (C-D) Ancillary GOS diet trials (2 and 3) showing that birds on the GOS diet (black) consistently achieved greater body weight at 35 da than birds provided with a calorie-matched control diet (blue). Significant differences are indicated by *p*-values above the diet pairs, with the corresponding cumulative feed conversion ratios (FCR) indicated in the bottom right hand side of each trial panel.

### *Lactobacillus* spp. distinguish microbial communities colonizing the ceca of broiler chickens on GOS-supplemented diets

Analysis of Trial 1 showed the *α*-diversity of the cecal microbiota was not significantly different between GOS and control diet cohorts sampled at 8, 15, 22 and 35 da (Inverse Simpson index: *p* ≥ 0.295; Shannon diversity, *p* ≥ 0.196; Supplementary Figure S1A and S1B). Community richness (Chao) was not significantly different throughout the trial (*p* ≥ 0.108, Supplementary Figure S1C). Communities of cecal bacteria colonizing birds on control or GOS diets could not be distinguished on the basis of Bray-Curtis dissimilarity indices at 8 (*p* ≥ 0.062) or 15 (*p* ≥ 0.087) da but could be distinguished at 22 (*p* = 0.013) and 35 da (*p* = 0.02, Supplementary Figure S1D).

The top 10 operational taxonomic units (OTU) with the greatest relative abundance are shown in Figure 2A for all the birds sampled at each time point. However, relatively few OTUs were discriminative between the control and GOS diets (Figure 2B). Two OTUs, OTU0006 and OTU0010 identified as *Lactobacillus* spp., were discriminatory of the different diets in the early rearing period up to 15 da (OTU0006, *p* ≤ 0.01; OTU0010, *p* ≤ 0.022). Organisms exhibiting 16S rRNA gene V4 region sequence identity with the differential OTUs identified by metagenomic analysis were isolated from MRS culture media. The genomic DNA sequences of these isolates were assembled from data generated from Illumina and PacBio platforms. Two representative isolates were designated *L. crispatus* DC21.1 (OTU0006) and *L. johnsonii* DC22.2 (OTU0010) based on whole genome alignments with type strains available in public nucleotide sequence databases.

**Figure 2.**
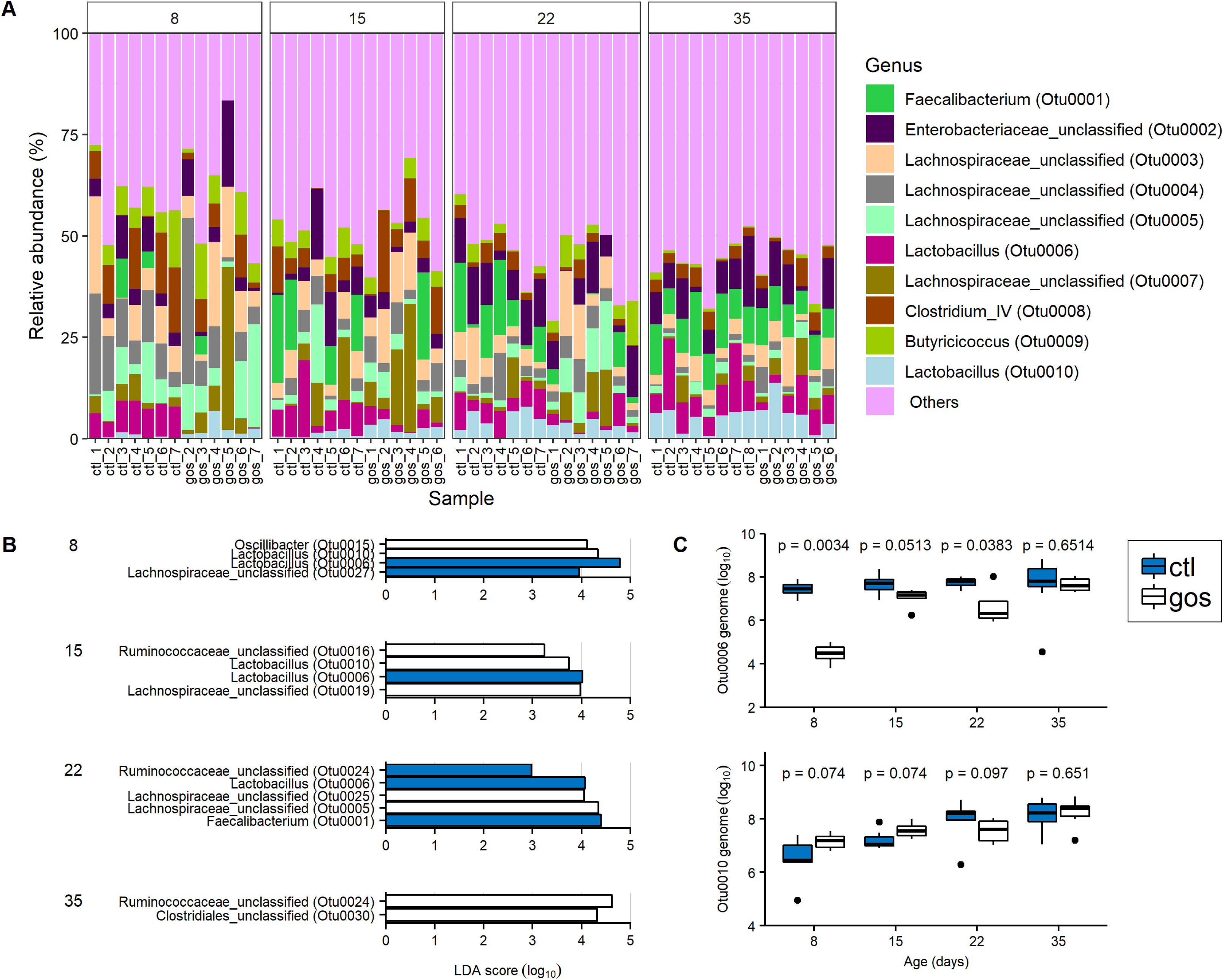
Metagenomic Analysis of GOS diet trial. (A) Stacked bar chart showing relative abundance to OTU level, comparing cecal contents from birds fed GOS diet with birds fed control diet. (B) Differential species identified in cecal contents from control diet (blue) and GOS diet (white) analyzed using Linear discriminant analysis effect size (LEfSe). (C) Box whisker plots of qPCR determinations of the cecal microbiota gene copy numbers of lactobacilli identified from birds fed on GOS (black) and control (blue) diets showing differential abundance. Significant differences are indicated by *p*-values above the sample pairs for n=7 with any outlying data points identified as spots.

Quantitative PCR assays were developed to measure the absolute abundance of *L. johnsonii* (OTU0010) and *L. crispatus* (OTU0006) within the gut microbiota. Oligonucleotide primers were designed on the *groEL* gene sequences as they have been frequently used to discriminate between *Bifidobacteria* strains with a high degree of similarity (26, 27). Once validated, using spiked cecal samples, the technique was used to enumerate *L. johnsonii* (OTU0010) and *L. crispatus* (OTU0006) within the cecal contents of control and GOS-fed birds. The genome copies of each isolate were measured throughout the rearing period. The results of these analyses confirmed the relative abundance estimates from metagenomic data in Trial 1 and demonstrated that the abundance of these two OTUs show positive and negative associations with birds fed the GOS diet compared to control diet (Figure 2C). Notably the abundance of *L. crispatus* (OTU0006) in the GOS-fed birds at 8 and 22 da was significantly lower than control birds (p < 0.05), and conversely *L. johnsonii* (OTU0010) was significantly greater at 8 and 15 da in the GOS-fed birds. Examination of the *L. crispatus*/*L. johnsonii* gene copy ratios from the cecal contents of birds over the trial revealed significant differences in the ratios (p < 0.05) observed between control and GOS-fed juvenile birds from 8 until 22 da.

### Characteristics of the *Lactobacillus* spp. distinguishing the cecal communities

Summaries of the functional gene contents of the *L. johnsonii* DC22.2 and *L. crispatus* DC21.1 isolates with respect to GOS utilization and host colonization are presented in Supplementary Table 1. *L. johnsonii* and *L. crispatus* have the capacity to colonize and compete in the host intestine with genes encoding multiple mucus binding proteins (28), fibronectin binding proteins (28), exopolysaccharide biosynthesis (29) and bile salt hydrolase (31). *L. johnsonii* carries *apf* gene (aggregation promoting factor) that encodes a cell surface protein that has been ascribed a role in cell adhesion (30). Whilst. *L. crispatus* contains the *cbsA* gene that encodes the structural protein that forms the S-layer (32, 33). *L. johnsonii* and *L. crispatus* both encode the bacteriocin helveticin J and multiple bacteriocin immunity factors.

There are ostensibly two pathways to utilize GOS that rely upon the cellular transporters LacS (lactose permease) or LacE/F (lactose phosphotransferase system), of which LacS permease appears capable of transporting GOS of DP2-6 but the LacE/F phosphotransferase is confined to DP2 lactose (34). The genome sequence of *L. johnsonii* DC22.2 suggests the isolate could be impaired in GOS utilization. The *lacS* permease gene has a stop codon at the seventeenth position, which would require that the protein be initiated from an internal AUG with the loss of the first 31 amino acids compared to the majority of database homologues. However, notable exceptions to this are *L. pasteuri* and *L. gallinarum* that respectively initiate translation at the corresponding position or 13 codons downstream. The syntenic position that harbors the lactose phosphotransferase encoding genes in the *L. johnsonii* poultry isolate FI9785 (35) features a deletion in *L. johnsonii* DC22.2 that preserves the *lacR* gene encoding the repressor but dispenses with all the functional components. In contrast, *L. crispatus* DC21.1 retains functional *lacS* and *lacA* genes to facilitate the use of GOS.

To assess the ability of the *L. crispatus* and *L. johnsonii* isolates to utilize GOS in axenic culture, the organisms were cultured in MRS basal medium containing DP2+ GOS in the absence of monosaccharides. Cultures were incubated for 72 h under anaerobic conditions at 37°C in basal medium with either DP2+ GOS (0.5 % v/v) or glucose (0.5 % w/v) as a positive control or sterile water instead of the carbon source as negative control (blank). Supplementary Figure S2 shows growth of the isolates and *Lactobacillus* type strains indicated by the OD_600_ measurements corrected for the negative control. *L. crispatus* DC21.1 utilizes GOS, showing increased growth over that recorded for *L. fermentum* ATCC 33323 in a parallel culture. Under these conditions *L. johnsonii* does not efficiently use GOS, which is consistent with the putative gene content but at odds with the differential abundance observed in the ceca of GOS-fed birds.

### In-feed GOS effects on gut architecture

Hematoxylin-Eosin (H&E) stains of ileal histological sections did not exhibit any significant differences in heterophil infiltration or inflammatory characteristics between control or GOS-fed birds at any time in the experiment (Figure S3A). The measurements of villus length and crypt depth indicate that GOS-fed juvenile birds at 15 da had longer villi (*p* = 0.05) and deeper crypts (*p* = 0.02) than birds on the control diet (Supplementary Table 2). However, these differences did not result in a difference in the villus/crypt ratio. At 22 and 35 da there were no significant morphometric differences recorded.

Goblet cell densities of villi from control and GOS-fed birds were evaluated from PAS-stained (neutral mucin producing) ileal sections (Supplementary Table 2). Greater densities of goblet cell were observed from GOS-fed birds sampled throughout Trial 1 with significant differences recorded at days 8 (*p* = 0.04), 15 (*p* = 0.002) and 22 (*p* = 0.04).

### In-feed GOS modulates host immune response

The immune responses were assessed in ileal and cecal tissues of Trial 1 by quantification of the relative expression of cytokine and chemokine genes representing the major inflammatory pathways of chickens (36).

The relative expression of the Treg-marker IL-10 and pro-inflammatory Th17 cell-associated IL-17A and IL-17F were profoundly modulated throughout the trial in both cecal and ileal tissues (Figure 3A; Supplementary Table 3). In cecal tissues cytokine expression in birds fed the GOS diet was marked by down-regulation of IL-10 (FC = 0.0001; *p* = 0.01) and IL-17F (FC = 0.1; *p* = 0.04) at 8 da and up-regulation of IL-17A at 8, 15 and 35 da (FC = >5; *p* < 0.01). Correspondingly, cytokine expression in ileal tissue was marked by down-regulation of IL-10 (FC = 0.0005; *p* = 0.002) and up-regulation of IL-17A (FC = 51; *p* = 0.002) at 8 da (Figure 3B, Supplementary Table 3), whilst IL-17F remained unchanged (*p* = 0.4). At 15 da the observed relative up-regulation of ileal IL-17A (FC = 4196, *p* = 0.003) and IL-17F (FC = 19; *p* = 0.004) is a consequence of a reduction in expression in the control birds, whilst expression in the GOS-fed birds remained similar to that at 8 da (Figure 3B). Similarly, in the ceca at 8 da down-regulation of IL-10 in the GOS-fed birds relative to control coincides with greater expression of IL-17A in birds on the GOS diet (Figure 3A).

**Figure 3.**
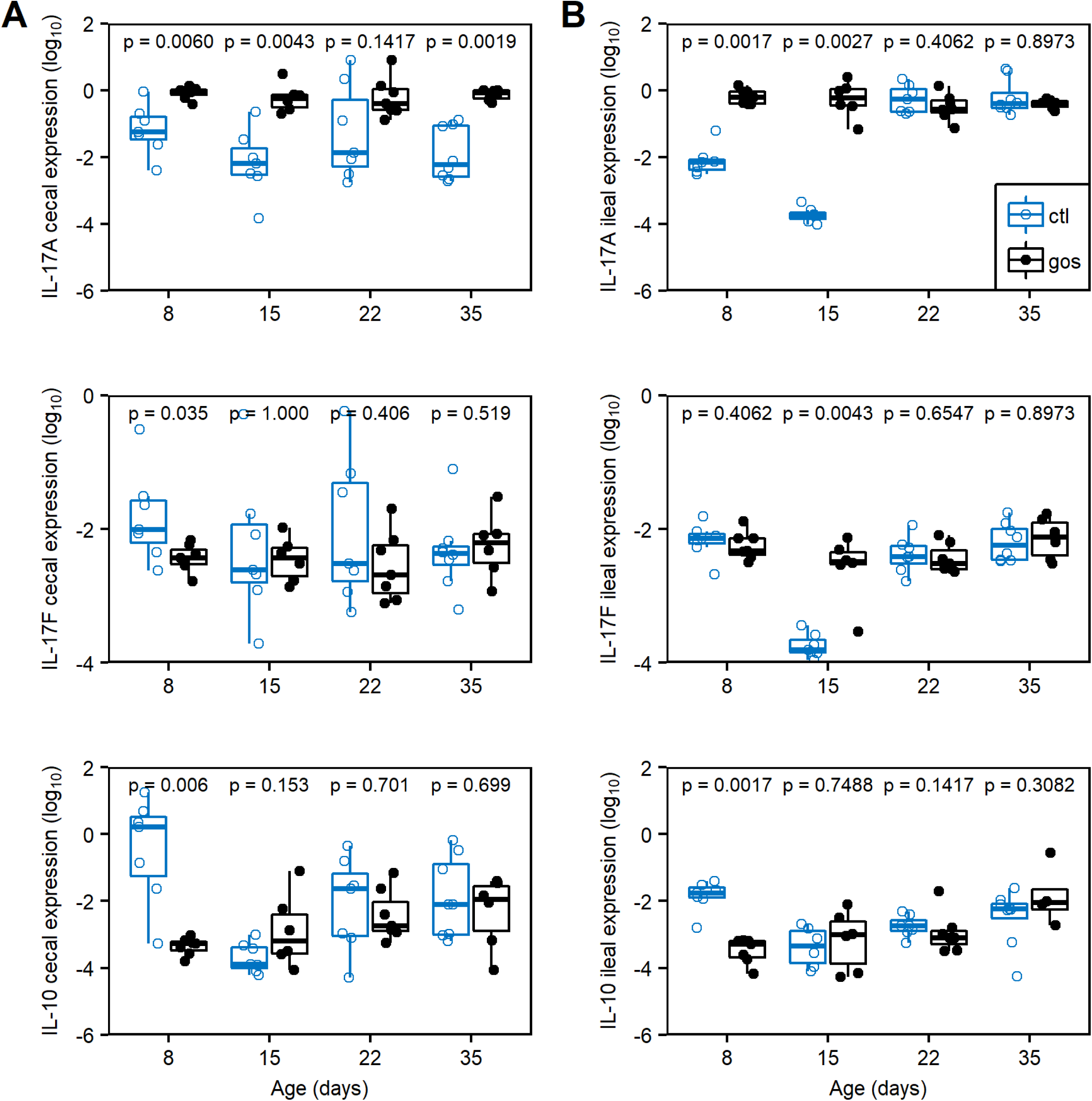
Changes in expression of IL-17A, IL-17F and IL-10 from cecal (A) and ileal (B) tissues. Relative gene expression was determined by quantitative RT-PCR, for GOS-fed birds (black) compared to expression from control diet (blue) fed birds. Expression of the gene of interest (GOI) relative to housekeeping gene (HG) is presented as box whisker plots of data from 7 birds determined from 3 technical replicates. The 2-Δ^Ct^ was determined for all samples such that positive value denotes increased expression and negative value denotes decreased expression. The ΔCt was calculated (Ct GOI – Ct HG) for each sample, significant differences between 2-ΔCt values of the control and GOS diet cohorts of birds are indicated by *p*-values above the sample pairs for n=7.

Expression of Th1-associated INF*γ* was up-regulated in the cecal tissues of GOS-fed birds at 15 da (FC = 11; *p* = 0.05) but not in older birds (Supplementary Table 3). In ileal tissues INF*γ* showed no differences between birds on control and GOS diets. Likewise, Th17 cell-associated IL-6 was upregulated in the ceca (FC = 1.5, *p* = 0.03) only at 15 da. IL-6 was also upregulated (FC = 1.5, *p* = 0.02) in the ileum at 35 da. The pro-inflammatory cytokine IL-4 was down-regulated in ileal tissue 15 da (FC = 0.02; *p* = 0.02). At 22 da in GOS fed birds pro-inflammatory chemokines ChCXCLi-1 and ChCXCLi-2 were reduced in the ceca (FC = 0.2; *p* < 0.03) and conversely increased in ileum (FC = 36, *p* < 0.02). These observations indicate that the GOS diet and/or concomitant shifts in the gut microbiota, do not drive induction of pro-inflammatory responses such as IL-1 *β* or Th1-associated IFN*γ* cytokines, whilst the reduction of IL-4, a marker for the Th2 pathway, was transient and limited to the ileum.

The increase of IL-17A transcription in the ceca of GOS-fed birds at 8 da coincides with reduced expression of the IL-10 (Figure S4D) and IL-17F transcripts (Figures S4E and S4F), suggesting that IL-17A and IL-17F are independently modulated. These data suggest that in-feed GOS induced elements of the Th17 immune response in the ceca, which coincides with greater relative abundance of *L. johnsonii* (OTU0010). Similarly, in the ileum at 8 da the in-feed inclusion of GOS resulted in higher levels of IL-17A and reduced levels of IL-10 transcripts (Figure S4A), whereas IL-17F transcripts were unaffected (Figure S4B and Figure S4C).

### Abundance of *L. johnsonii* in the cecal lumen positively correlates with bird growth performance

Correlations between abundances of the *Lactobacillus* isolates with bird weight were analyzed by combining the data for 35 da birds from trials that represent a range of performance outcomes (Trials 1,4 and 5). These data show a clear positive relationship between body mass and the *L. johnsonii* genome copy number per gram of cecal content determined by qPCR (rPearson = 0.888; *p* < 0.001; Figure 4A). The abundances of *L. crispatus* and *L. johnsonii* were further analyzed for any relationship with the expression of IL-17A, IL-17F and IL-10. Figure 4B shows the correlation noted with the expression of IL-17A and *L. johnsonii* genome copy number (rPearson = 0.487; *p* = 0.004). No significant correlation of IL-17A with *L. crispatus* was evident. These results taken together strongly suggest that *L. johnsonii* acts as a key species promoted by GOS to improve growth performance and prime a Th17 immune response.

**Figure 4.**
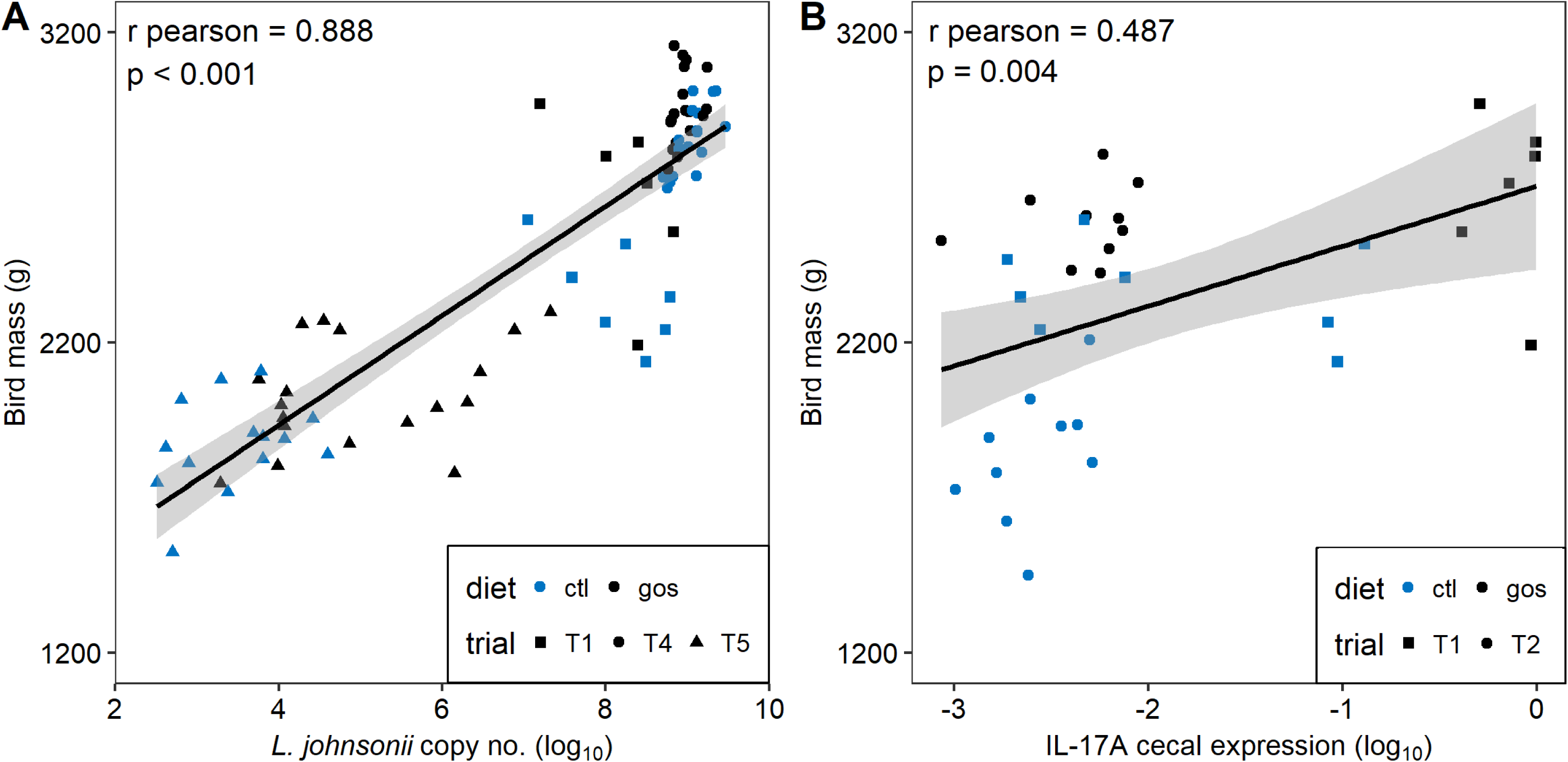
Correlation of growth performance to *L. johnsonii* abundance and IL-17A expression. Panel A shows the Pearson correlation of bird mass at 35 da against *L. johnsonii* gene copy number gene copy number per g cecal content. Panel B shows the Pearson correlation of bird mass at 35 da against cecal IL-17A gene expression.

### Modulation of cecal lactobacilli and bird growth performance

To examine if shifts in the cecal abundance of lactobacilli of juvenile broiler chickens can modify growth performance, we administered either *L. crispatus* DC21.1 or *L. johnsonii* DC22.2 (8 log_10_ CFU) by cloacal gavage to chicks at 6 da. Cloacal gavage has the advantage of allowing cecal colonization without the impact of upper intestinal transit and accompanying losses in the effective dose of the colonizing bacteria (37). Figure 5A shows marked shifts in the cecal abundance ratios of *L. crispatus*/*L. johnsonii* (competitive indices calculated from genome copy numbers per gram of cecal content) at 35 da in favor of the *Lactobacillus* spp. administered compared to the non-treated controls for birds on control or GOS diets. Extreme differences in the cecal abundance of *L. johnsonii* corresponded with disparate differences in the weights of mature birds at 35 da. The mean body weights of the birds with low cecal *L. johnsonii* abundance (administered with competitive *L. crispatus*) fed on control diet were 2.0 ±0.2 kg compared with 2.8 ±0.1 kg for the birds with high cecal *L. johnsonii* abundance (*p* < 0.001). Figures 5B and 5C show the respective correlations between body mass and *L. johnsonii* genome copy number from the cecal microbiota of birds administered with either *L. johnsonii* DC22.2 (rPearson = 0.353; *p* = 0.038) or *L. crispatus* DC21.1 (rPearson = 0.511; *p* = 0.003). These data confirm the relationship of *L. johnsonii* DC22.2 with growth performance and that *L. crispatus* DC21.1 is a competitor in the development of the cecal microbiota. Early shifts in the juvenile microbiota have a profound effect on the weights of market ready broiler chickens. Figure 5C shows the impact of dietary GOS is greatest on the weaker performing birds administered with *L. crispatus* with increases in body weight and increases in cecal *L. johnsonii* abundance due to expansion of the niche available to the resident bacteria.

**Figure 5.**
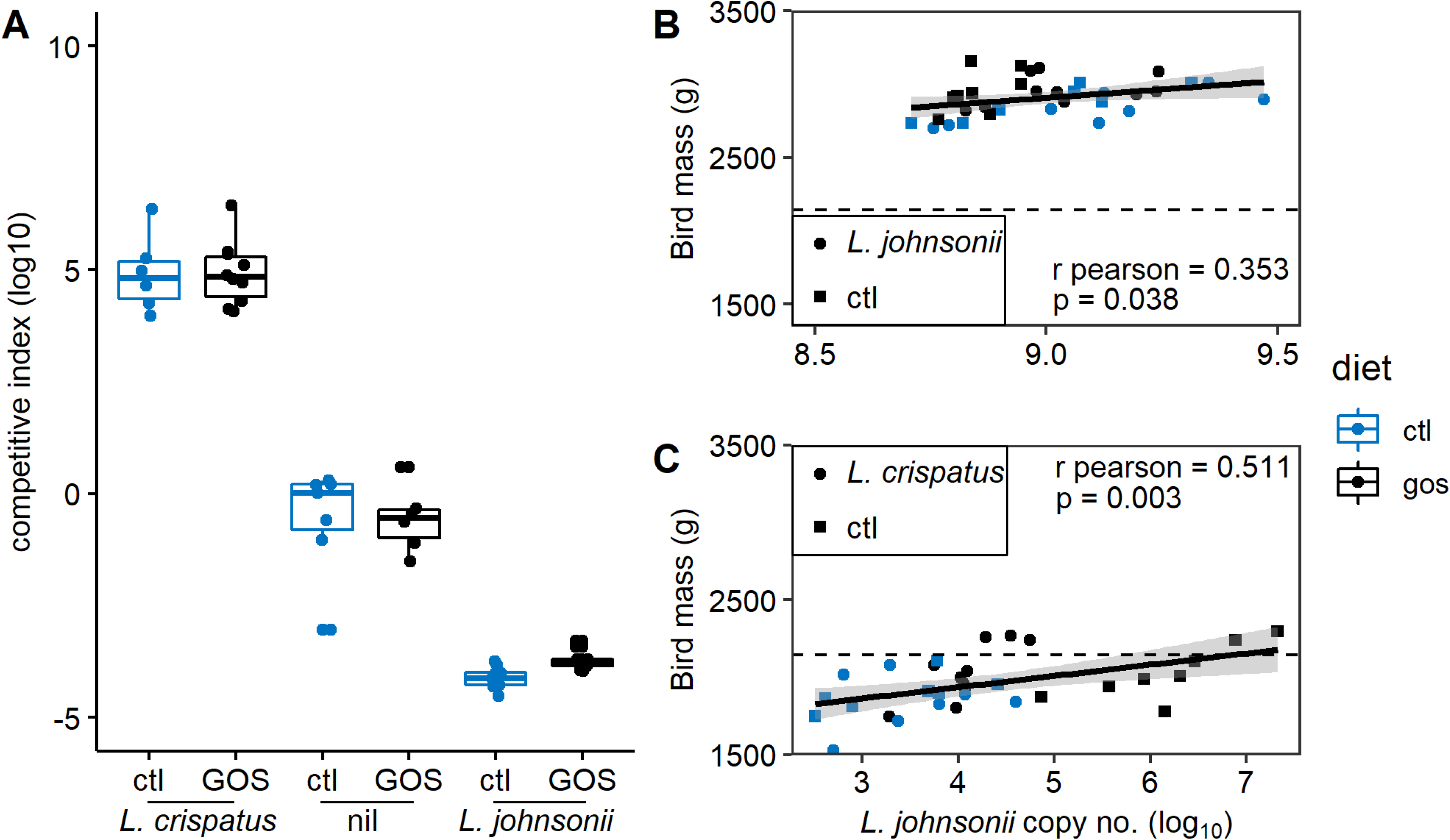
Modulation of the juvenile cecal microbiota. Panel A shows the competitive indices (*L. crispatus*/*L. johnsonii* ratios) at 35 da of broiler chickens either non-treated or administered with either 8 log_10_ CFU *L. crispatus* DC21.1 or *L. johnsonii* DC22.2 by cloacal gavage at 6 da. Competitive indices were calculated from *L. crispatus* or *L. johnsonii* genome copy numbers determined by qPCR from the cecal microbiota. Panel B shows the Pearson correlation of bird mass at 35 da against *L. johnsonii* gene copy number per g cecal content in birds administered *L. johnsonii*. Panel C shows the Pearson correlation of bird mass at 35 da against *L. johnsonii* gene copy number per g cecal content in birds administered *L. crispatus*. The dashed horizontal line represents the male Ross 308 Performance Objective (Aviagen 2014).

## DISCUSSION

Improvements in the growth performance together with improved health are key goals in broiler chicken production. The inclusion of galacto-oligosaccharides in broiler feed resulted in enhanced growth rate relative to chickens on a calorie-matched control diet. Prebiotic galacto-oligosaccharides have previously been reported to improve the performance, intestinal architecture and stimulate intestinal defenses of neonatal pigs (38). However, a GOS supplemented diet fed to chickens increased fecal populations of bifidobacteria and lactobacilli but did not improve zootechnical performance (19). In contrast, we observed a significant improvement in performance of the GOS-fed broiler chickens that was also accompanied by changes in the intestinal microbiota. Differences in the abundance of specific members of the *Lactobacillus* genus in the cecal microbiota of juvenile birds on control and GOS diets were observed. *Lactobacillus* isolates were recovered from the cecal contents of the birds, and specific isolates identified that shared sequence identity with the OTUs displaying differential abundance in the microbiota of birds consuming either GOS or control diets. Whole genome sequence alignments allowed the identification of *L. johnsonii* isolate DC22.2 prevalent in the ceca of GOS-fed birds and a *L. crispatus* isolate DC21.1 exhibiting greater abundance in control birds.

Lactobacilli need to import GOS (degree of polymerization 2–6) and lactose since the enzymes required to breakdown the substrate are generally cell bound. Two principal pathways accomplish import: LacS lactose permease or the LacE/F phosphotransferase system, where the latter appears to be restricted to DP2 lactose (34). The dependence of LacS for the utilization DP>2 GOS was first established in *L. acidophilus* (39). Intestinal *Lactobacillus* species that form the “acidophilus complex” include *L. crispatus*, *L. johnsonii*, and *L. helveticus*, which show conservation of the *gal*-*lac* clusters (39). Whilst the *lacS* gene is often present, the co-presence of *lacS* and *lacA* (encoding β-galactosidase) appears to be associated with their ability to utilize GOS. *L. johnsonii* isolates exhibit host specific differences, where human and porcine sources frequently possess the *lacS* and *lacA* genes but the poultry isolate FI9785 appears deficient due to genome rearrangements (35, 40). However, isolate FI9785 does encode orthologues of *lacE/F*. Converse to this *L. johnsonii* DC22.2 has retained *lacS* and *lacA* but has lost the *lacE/F* genes. Paradoxically in our experiments *L. johnsonii* DC22.2 does not efficiently utilize GOS *in vitro* but represents a greater differentially abundant component of the cecal microbiota of GOS-fed birds that exhibit improved zootechnical performance. In contrast, *L. crispatus* DC21.1 can utilize GOS but this did not provide a competitive advantage in GOS-fed chickens, rather the reverse appears to be true. It seems unlikely that *L. johnsonii* DC22.2 can compete for the GOS substrate directly with *L. crispatus* DC21.1, which suggests that *L. johnsonii* DC22.2 is benefiting from the metabolic capability of another member of the cecal microbiota. The presence of GOS provides the trophic selection for members of the cecal community required to support autochthonous *L. johnsonii*. Indeed, a hallmark of the acidophilus complex gene contents is the absence of the biosynthetic pathways necessary to produce essential nutrients such as amino acids, purine nucleotides and cofactors, and therefore a reliance on effectively importing nutrients generated by the intestinal milieu (41; 42). Prebiotic selection *in situ* may well be a more reliable way of achieving a beneficial microbiota than directly providing dietary probiotics as the response will be congruent with the metabolic capabilities of the resident community. Prevailing environmental conditions may alter the content and composition of the intestinal microbiota, and therefore the outcomes of prebiotic selection. For example, members of the *Bifidobacterium*, *Christensenella* and *Lactobacillus* genera have been reported to feature in the intestinal communities of chickens on GOS diets (19, 43).

Regardless of the route of succession, increased levels of *L. johnsonii* in GOS fed birds correlate with improved performance. *L. johnsonii* is an established probiotic with a variety of reported effects when administered to humans and animals. For example, *L. johnsoni* isolate N6.2 has been shown to have immunomodulatory effects in animal experiments and to protect diabetes prone rats from developing the disease (44, 45). In another study *L. johnsonii* was shown to attenuate respiratory viral infection via metabolic reprogramming and immune cell modulation (46). *L. johnsonii* LB1 expresses a bile salt hydrolase active against tauro-beta-muricholic acid (T-*β*-MCA), a critical mediator of farnesoid X receptor (FXR) signaling that is important in maintaining metabolic homeostasis (47). In broiler chickens, administration of a *L. johnsonii* isolate has been reported to improve growth performance (48). Subsequently it was reported that meat from *L. johnsonii* treated birds had higher nutritional value and the birds showed resistance to the development of necrotic enteritis (49, 50). Similar results were obtained by administering *L. johnsonii* LB1 to piglets to improve performance and reduce diarrhea (51). The administration of *L. johnsonii* FI9785 to broiler chickens has also been reported to reduce colonization by *Clostridium perfringens*, *E. coli* O78:K80 and *Campylobacter jejuni* that have a significant impact on poultry production (52, 53). *Lactobacillus crispatus* is also a recognized probiotic, but better known for its activity against recurrent urinary infections, bacterial vaginosis, and vaginal candidiasis (54). However, it should be noted that *L. crispatus* is commonly reported as a major constituent of the chicken microbiome (43, 52).

Shifts in the *Lactobacillus* spp. in response to GOS are accompanied by changes in the expression of cytokines and chemokines that have the potential to prime innate intestinal immune systems and enhance pathogen resistance. However, unchecked low-grade pro-inflammatory responses can cause tissue damage and inefficient feed conversion (55). Lactic acid, for example, is a by-product of glycolytic pathway of immune cells that can affect local T cell immunity by inhibiting T cell motility, and inducing the change of CD4+ cells to a Th17 pro-inflammatory T cell subset, which leads to IL-17 production and chronic inflammation (56). However, lactic acid is also the homofermentative product of lactic acid bacteria such as *L. johnsonii*, the action of which in the gut has recently been reported to promote the expansion of intestinal stem cells, Paneth cells and goblet cells (57). Coincident with the increased abundance of *L. johnsonii* we observed significantly greater ileal goblet cell densities from juvenile birds on the GOS diet. We also observed an increase in IL-17A and decreases in IL-10 and IL-17F gene expression in juvenile birds on GOS diet. IL-17A has been proposed to promote the maintenance of intestinal epithelial cell integrity based on observations that IL-17A inhibition exacerbates colitis in a mouse model that leads to weakening of the intestinal epithelial barrier (58). In contrast, IL-17F knockout mice are reported to be protected against chemically induced colitis, whereas IL-17A knockout mice remain sensitive (59). Moreover, IL-17F deficient mice increased the colonic abundance of *Clostridium* cluster *XIVa* that promote beneficial regulatory T cells, and the expression of β-defensins 1 and 4 (59). Extrapolating from these data we propose that the increased expression of IL-17A we observe at the expense of IL-17F in GOS-fed chickens will promote gut health, a prerequisite for improved commercial production.

The induction of host Th17 responses in ileal and cecal tissues by an indigenous symbiont is reminiscent of the Th17 stimulation brought about by adherent segmented filamentous bacterium (SFB) described in mice (60, 61). The tight adherence of SFB to epithelial cells was observed to accelerate postnatal maturation of intestinal mucosal immunity by triggering a Th17 response (62, 63). The observation of an intestinal Th17 response to tightly adherent symbiotes was extended to the human symbiont *Bifidobacterium adolescentis* in mice (64). However, the response arose through a distinctly different transcriptional program to that observed for SFB, suggesting that intestinal Th17 responses are maintained by parallel sensor and signaling pathways. The up-regulation of IL-17A in juvenile chickens fed on a GOS diet may also support gut immune maturation through Th17 cell stimulation. In chickens, lactobacilli are recognized adherent species to the intestinal tract epithelium from crop to ceca. Besides the cell adhesion factors we note in our isolates, for *L. johnsonii* additional roles have been proposed for surface-associated GroEL, elongation factor Tu and lipoteichoic acid (65–68).

Juvenile chickens have been reported to exhibit a transient IL-17 induction during the development of the natural microbiota (69). The study suggested that in the absence of IL-22, pro-inflammatory Th17 induction did not result in intestinal tissue damage. It is possible that IL-17 has a role in the co-development of the microbiota and innate immunity in chickens, which is consistent with our findings that up-regulation of IL-17A did not cause lamina propria inflammation. Crhanova et al. (69) also reported the outcome of *Salmonella* Enteritidis challenge of chickens shifts from a Th1 response (induction of interferon-γ and nitric oxide synthase) at 1-4 da, to a Th17 response at 16 da (induction of IL-17). They conclude that a mature Th17 subset of helper T cells produced IL-17 and IL-22, which confers resistance to *S*. Enteritidis infection and damage in older birds.

### Conclusion

We have demonstrated an increase in growth rate of broiler chickens in response to the dietary supplementation with the prebiotic GOS. Juvenile chickens on GOS starter feed exhibit differences in the cecal abundance of key species of Lactobacillus compared to control feed. Differences in the cecal microbiota in early development correlate with the composition of the mature cecal microbiota and performance outcomes. The provision of dietary GOS increased the density of goblet cells populating ileal villi in the developing chicken gut. Goblet cell increases were accompanied by significant differences in the villus height and crypt depth at 15 da, a period when transitions in the development of the chicken microbiota from a juvenile to a mature composition are observed (70–72). Using molecular methods, we have demonstrated a significant correlation between the market weight of chickens at 35 da and the cecal abundance of a specific *L. johnsonii* isolate identified as differentially abundant in the juvenile microbiota of GOS-fed birds. We further observed a significant correlation between IL-17A gene expression and the abundance of *L. johnsonii*. Prebiotic GOS diet increased IL-17A gene expression and decreased the expression of IL-10 and IL-17F in ileal and cecal tissues of developing chickens. IL-17A has been proposed to have beneficial effect on gut health, whilst IL-17F expression a role in the development of colitis (56, 57). The correlation between increased IL-17A expression and *L. johnsonii* abundance parallels the intestinal induction of IL-17A observed in response to tightly adherent symbiotic species (60–64). *L. johnsonii* is an established probiotic that has been demonstrated to have beneficial effects when applied in poultry production (48–53). Several modes of action have been proposed for probiotic strains of *L. johnsonii* but underlying these is the multi-modal ability of the species to affect epithelial gut cell adherence (65–68), which we propose will induce the expression of IL-17A. By taking a system-wide approach we have, for the first time, established mechanistic links between prebiotic selection of an autochthonous synbiotic species, increased IL-17A expression and the development of the gut in healthy animals.

## MATERIALS AND METHODS

### Ethical approval

Experiments involving the use of birds were subjected to approval process under national guidelines by the United Kingdom Home Office. Work on this project was approved under United Kingdom Government Home Office Project Licensing ASPA 86. All project licenses are reviewed internally by the University Ethics Committee prior to submission to the Home Office. This includes the scrutiny of animal welfare, ethics and handling.

### GOS Diet Trial 1

#### Experimental birds

Commercial male Ross 308 broiler chicks (*n* = 70) were obtained as hatchlings (PD Hook, Oxfordshire, UK). Birds were housed in a controlled environment under strict conditions of biosecurity. Temperatures were as outlined in the Code of Practice for the Housing and Care of Animals Bred, Supplied or Used for Scientific Purposes. Birds were provided feed and water *ad libitum*. Feeds were formulated on a least cost basis and to meet the requirements set out in the Ross 308 Broiler: Nutrition Specifications 2014 (Aviagen, UK), and prepared by Target Feeds Ltd (Shropshire, UK). The diet regime was as follows: the control diet group was sustained on a wheat-based diet provided as a starter crumb for 0-10 days of age (da), grower pellets for 11-24 da and finisher pellets for 25-35 da. The starter diet contained wheat (59.9% w/w), soya meal (32.5% w/w), soyabean oil (3.65% w/w), limestone (0.6% (w/w), calcium phosphate (1.59% w/w), sodium bicarbonate (0.27% w/w), the enzymes phytase and xylanase (dosed according to the manufacturer’s instructions; DSM Nutritional Products Ltd., PO Box 2676 CH-4002 Basel, CH) and a vitamin mix containing salt, lysine HCl, DL-methionine and threonine. The grower and finisher diets increased the wheat content at the expense of soya meal by 2 and 5% w/w respectively. GOS was provided as Nutrabiotic® GOS (Dairy Crest Ltd, Davidstow, Cornwall, UK). Galacto-oligosaccharide preparations contain a mixture of monosaccharides (glucose and galactose) and oligosaccharides (DP2 – DP8). The disaccharides lactose, a reactant in the manufacture of galacto-oligosaccharides, is not a galacto-oligosaccharide; all other disaccharides and longer (DP3+) oligosaccharides are considered to be galacto-oligosaccharides and non-digestible. The starter feed was supplemented with 3.37% w/w GOS and isocaloric adjustments made in the wheat (54.0% w/w) and soybean oil (4.88% w/w) contents. The grower and finisher feeds contained 1.685% GOS with the respective adjusted wheat contents of 57.7% (w/w) and 63.3% (w/w), and soybean oil contents of 6.14% (w/w) and 6.22% (w/w). The final feeds were isocaloric (ME including enzyme contribution), contained the same crude protein levels and Degussa poultry digestible amino acid values (lysine, methionine, methionine + cysteine, threonine, tryptophan, isoleucine, valine, histidine and arginine).

Chickens were euthanized by either exposure to rising CO_2_ gas or parenteral barbiturate overdose followed by cervical dislocation according to Schedule 1 of the UK Animals (Scientific Procedures) Act. The birds were weighed before tissue and intestinal content were sampled post-mortem. Ileal tissues were sectioned from approximately 3 cm distal to Meckel’s diverticulum and cecal tissues collected from the distal tips of the ceca. Intestinal tissues were immediately frozen in liquid nitrogen for subsequent RNA isolation or preserved in 10% (w/v) neutral buffered formalin (Fisher Scientific, Loughborough, UK) for histological assessment. Intestinal contents were collected and stored at −80°C until DNA isolation.

#### Zootechnical performance

All birds were weighed, and the feed consumed recorded at least weekly from the start of the experiment until the end at 35 da. Feed conversion ratios (FCR) were calculated as a ratio of feed consumed to the cumulative weight of the birds.

#### Cloacal gavage

Axenic suspensions of either *L. crispatus* DC21.1 or *L. johnsonii* DC22.2 containing 8 log_10_ CFU in 0.1 ml MRD (Oxoid, Basingstoke, UK) were administered to 6 da chicks by cloacal gavage performed using a blunt narrow-nosed syringe. The birds were fed either the control diets throughout or the GOS diets to 22 da and then control to end of the trial at 35 da.

### Ancillary GOS Diet Trials

Further GOS diet trials were carried out using the starter and grower feed formulations detailed as above. However, the GOS treatment and control birds in these trials were fed the control finisher diet 25-35 da. The organization of Trial 2 was as Trial 1, whilst for Trials 3-5 the birds were wing-tagged and housed in treatment groups in pens instead of individual caging.

### Isolation and enumeration of coliforms and lactic acid bacteria

Cecal contents from each individual bird were serial diluted in MRD (Oxoid) and spread (0.1 ml) onto the surface of MacConkey No 3 or MRS (Oxoid) plates. The MacConkey plates were incubated at 37 °C, for 24 h. The MRS plates were incubated under anaerobic conditions for 48 h at 37 °C. The numbers of coliforms and lactic acid bacteria colonies were recorded and examples of distinct, well isolated colonies from the MRS plates were sub-cultured for identification and storage at −80°C.

### Identification of lactic acid bacteria

Multiple isolates from MRS plates were examined by microscopy using the Gram stain. Genomic DNAs were prepared from selected isolates showing different cell and colony morphologies using GenElute™ Bacterial Genomic DNA Kit (Sigma Aldridge, Gillingham, UK). Identification to presumptive species level was carried out by performing PCR amplification of 16S rRNA gene sequences using primers TPU1 and RTU8 (73; Supplementary Table 4) and DNA sequencing of the products following clean-up (Wizard SV Gel and PCR Clean-Up System, Promega, Southampton, Hampshire, UK) using dye terminator chemistry (Eurofins, Ebersberg, Germany). The 16S rRNA gene V4 region sequences were matched to the OTU clusters outputted from microbiome analysis. The genome sequences of *L. crispatus* DC21.1 and *L. johnsonii* DC22.2 were assembled using CLC Genomics Workbench 10.0.1 (Qiagen, Aarhus, Denmark) using a combination of data generated from the Illumina MiSeq and PacBio RSII platforms. The *L. crispatus* DC21.1 and *L. johnsonii* DC22.2 cultures were deposited at National Collection of Industrial Food and Marine Bacteria (NCIMB) under the respective accession numbers 42771 and 42772.

### *In vitro* growth of Lactobacillus on galacto-oligosaccharides

To determine the ability of *L. crispatus* DC21.1 and *L. johnsonii* DC22.2 to utilize GOS *in vitro*, a purified Nutrabiotic® GOS (74% w/w dry matter) containing a DP2+-lactose fraction was prepared using HPLC, with an Imtakt Unison UK-Amino (aminopropyl stationary phase) column (ARC Sciences, Oakham, UK) with acetonitrile-water mobile phase, to remove monomeric sugars (glucose and galactose) and lactose (IPOS Ltd, Huddersfield, UK). A reduced carbon source medium based on MRS broth with the omission of glucose was prepared as a basal medium (pH 6.7). One litre of the medium contained: 10 g tryptone (Oxoid), 5 g yeast extract (Oxoid),10 g Lab-Lemco powder (Oxoid), 1 ml Sorbitan mono-oleate (Tween 80), 2 g di-potassium hydrogen phosphate, 0.5 g sodium acetate 3H_2_O, 2 g di-ammonium hydrogen citrate, 0.2 g magnesium sulfate 7H_2_O, and 0.05 g manganese sulfate 4H_2_O (from Fisher Scientific unless otherwise stated). Each experiment was carried out using the basal medium with addition of DP2+-lactose GOS (0.5 % v/v), together with a positive control, containing glucose (0.5 % w/v) and a negative control with sterile water instead of the carbon source. The bacterial cultures were grown on MRS plates and suspended in the modified MRS medium to a density of 8 log_10_ CFU/ml (OD_600_ of approximately 1.5). The suspension was diluted 1 in 100 into the growth medium. The assay was carried out in triplicate with 3 technical replicates per biological replicate together with a set of un-inoculated negative controls as blanks (0.2 ml in microtitre plates). The plates were covered and incubated at 37°C for 72 h, under anaerobic conditions with shaking. The OD_600_ obtained from growth on the basal medium, without the addition of carbon source, was subtracted from the value of the growth on the selected carbon source.

### Histology

Tissue samples fixed in a 10% formalin solution were dehydrated through a series of alcohol solutions, cleared in xylene, and embedded in paraffin wax (Microtechnical Services Ltd, Exeter, UK). Sections (3 to 5 *µ*m thick) were prepared and stained with either modified hematoxylin and eosin (H&E) or periodic acid–Schiff (PAS) using standard protocols. After staining, the slides were scanned by NanoZoomer Digital Pathology System (Hamamatsu, Welwyn Garden City, UK). Measurements of villus height and crypt depth were made using the NanoZoomer Digital Pathology Image Program (Hamamatsu) of 10 well-oriented villi scanned at 40X resolution for each tissue sample. Villus height was measured from the tip of the villus to the crypt opening and the associate crypt depth was measured from the base of the crypt to the level of the crypt opening. The ratio of villus height to relative crypt depth (v:c ratio) were calculated from these measurements. Goblet cells were enumerated from ileal sections stained with PAS. Measurements of 10 well-oriented villi per tissue sample of 3 or 4 birds per diet group at each sampling time were analyzed.

### RNA isolation and RT-qPCR of the cytokines and chemokines

RNAs were isolated from ceca and ileum tissue biopsies using NucleoSpin RNA purification kit (Macherey-Nagel, GmbH & co. KG, Düren, Germany) according to the manufacturer’s protocol with the following modifications. Tissue samples were homogenized with the kit Lysis buffer and 2.8 mm ceramic beads (MO BIO Laboratories Inc., Carlsbad, USA) using TissueLyser II (Qiagen, Hilden, Germany). Subsequently total RNA was extracted as described in the protocol with a DNase I treatment step as per the manufacturer’s instructions. Purified RNAs were eluted in nuclease free water, validated for quality and quantity using UV spectrophotometry (Nanodrop ND-1000, Labtech International Ltd, Uckfield, UK), and stored long term at −80°C. RNAs with OD _260/280_ ratio between 1.9 and 2.1 were deemed high quality, the ratios were found with a mean of 2.12 plus minus 0.01. Reverse Transcription was performed with 1 *µ*g of RNA, SuperScript II (Invitrogen Life Technologies, Carlsbad, USA.) and random hexamers as described previously (70).

Quantitative PCR was performed with cDNA templates derived from 4 ng of total RNA in triplicate using SYBR Green Master mix (Applied Biosystems, ThermoFisher Scientific, UK). The RNA level of expression was determined by qPCR using the Roche Diagnostics LightCycler 480 (Hoffmann La Roche AG, Switzerland). The primers sequence for GAPDH, INF*γ*, IL-1 *β*, IL-4, IL-6, IL-10, IL-17A, IL-17F, ChCXCLi1 and ChCXCLi2 (74,75,76) are presented in Supplementary Table 4. Cytokines and chemokines transcripts fold change were calculated according to the manufacturer’s recommendation using the 2^−ΔΔCt^ method (77). The means of triplicate Ct values were used for analysis, where target genes Ct values were normalized to those of the housekeeping gene glyceraldehyde 3-phosphate dehydrogenase (GAPDH).

### Microbiome analysis

DNA isolation from cecal content DNA was isolated from cecal content using the MoBio PowerSoil kit (now QIAGEN Ltd, Manchester, UK) according to the manufacturer’s instructions. For microbiome analysis the V4 regions of the bacterial 16S rRNA genes were PCR amplified using the primers 515f and 806r (Supplementary Table 4) (78). Amplicons were then sequenced on the Illumina MiSeq platform using 2 × 250 bp cycles. The 16S rRNA gene sequences were quality filtered and clustered into OTUs in Mothur (79,80) using the Schloss lab. MiSeq SOP (https://www.mothur.org/wiki/MiSeq_SOP, accessed 2018-10-05; 81). Batch files of Mothur commands used in this study are available at https://github.com/PJRichards/Richards_GOS_broiler. Raw sequence data are deposited in the NCBI database within the Bioproject PRJNA380214. Post-processing rarefaction curves were plotted to assess sampling effort (Figure S5).

### Quantitative PCR enumeration of lactobacilli

Quantitative PCR protocols to enumerate *L. crispatus* DC21.1 and *L. johnsonii* DC22.2 from intestinal contents were developed by designing primers specific for the *groEL* gene sequences of these bacterial strains (Supplementary Table 4). Real-time qPCR quantification of *L. crispatus* DC21.1 and *L. johnsonii* DC22.2 was performed with 1µl of cecal content DNA (15-150 ng) using SYBR Green Master mix with the Roche Diagnostics LightCycler 480. The amplification conditions were denaturation at 95°C for 5 minutes followed by 45 cycles of 95°C for 15 seconds denaturation and 60°C for 1 minute annealing. The fluorescence signals were measured at the end of each annealing step. Melting curves were generated by heating the samples from 65°C to 97°C at a ramp rate of 0.11°C per second. The data obtained was plotted against a standard curve generated with five-fold serial diluted target bacterial DNAs. Genome copy number of target bacteria in each dilution was calculated based on its genome length and applied DNA quantity with the assumption of the mean molecular mass of one base pair as 650 Daltons. The method was validated first with DNA extracts from pure cultures of known CFU, and then by spiking chicken cecal samples with increasing concentrations of target cells. The data was calculated as genome copy number per microliter of DNA applied in the PCR reaction and converted into genome copy number per gram of cecal content based on the mass of cecal material and elution volume applied for DNA extraction.

### Data and Statistical Analysis

For the microbiota β-diversity analysis Bray Curtis distances were tested for significance using AMOVA implemented within Mothur (79). Linear discriminant analysis effect size (LEfSe) was used to identify differentially abundant OTUs (82) and implemented within Mothur (79). Data processing and ordination were performed using R (R Development Core Team, 2013. R: A language and environment for statistical computing. R Foundation for Statistical Computing, Vienna, Austria. ISBN 3-00051-07-0; URL http://www.R-project.org; 83). With the exception of Figure S2 all figures were drawn using R version 3.5.3 (2019-03-11) in Rstudio 1.1 (84,85). R scripts used to draw figures are available at: https://github.com/PJRichards/Richards_GOS_broiler. Significance tests of heterophil counts, villus and crypt measurements were performed using single-factor ANOVA with *p* < 0.05 used as the level significance. Non-normally distributed gene expression data were compared using the non-parametric Kruskal-Wallis test.

### Data Availability

The genome DNA sequences of *L. crispatus* DC21.1 appear in the NCBI database under the accession numbers CP039266-CP039267. The genome DNA sequences *L. johnsonii* DC22.2 appear in the NCBI database under the accession numbers CP039261-CP039265. Raw DNA sequence data and metadata in support of 16S rRNA metagenomic analysis appear in the NCBI database within Bioproject PRJNA380214.

## ACKNOWLEDGEMENTS

The authors acknowledge research funding from Dairy Crest Ltd. NMF is a consultant to Dairy Crest Ltd.

---

**Figure S1.**
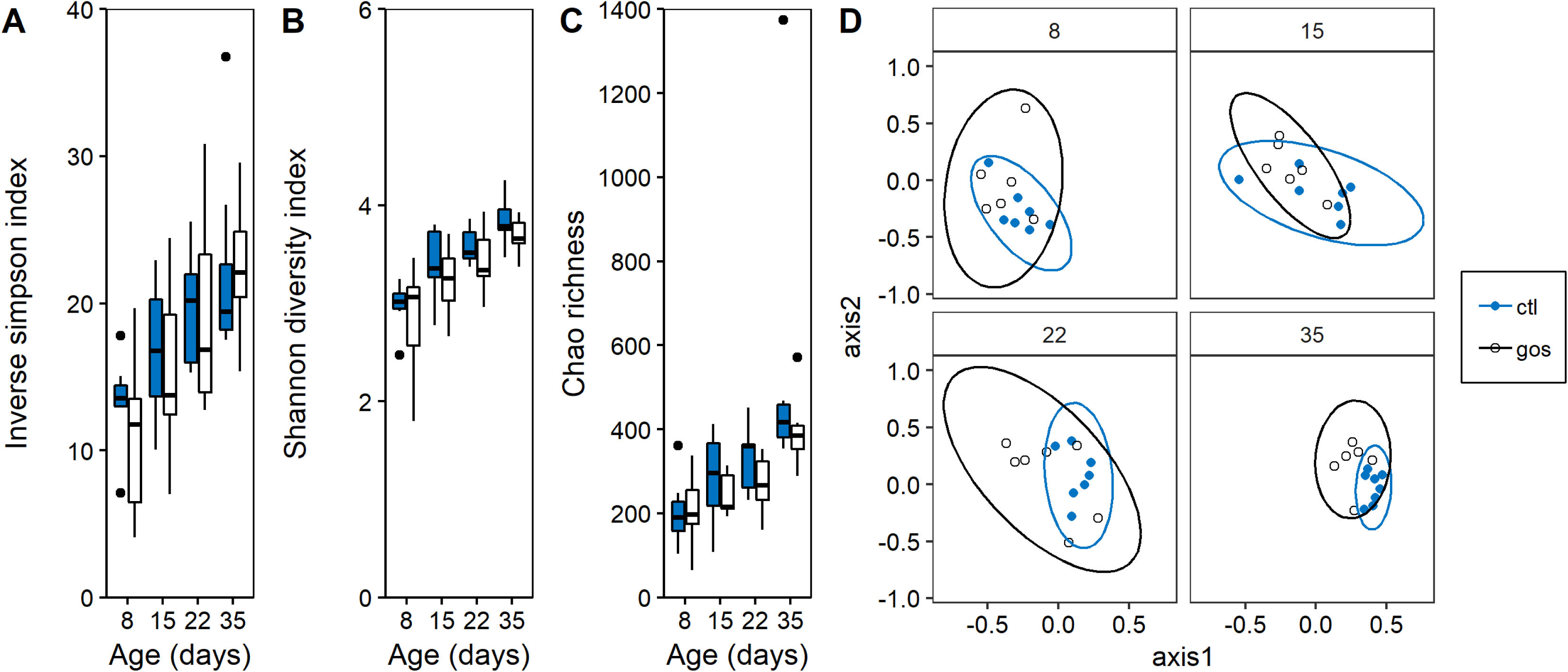
Diversity of the cecal microbiota on GOS and control diets. Panels A-C show indices of α–diversity of cecal bacterial populations for broiler chickens on GOS (black) and control (blue) diets sampled through the rearing period (35 da). Panel D shows NMDS plots of Bray-Curtiss dissimilarity indices for all birds at all sample days. Sampling days 22 and 35 show significant differences in the cecal microbial communities (*p* ≤ 0.02).

**Figure S2.**
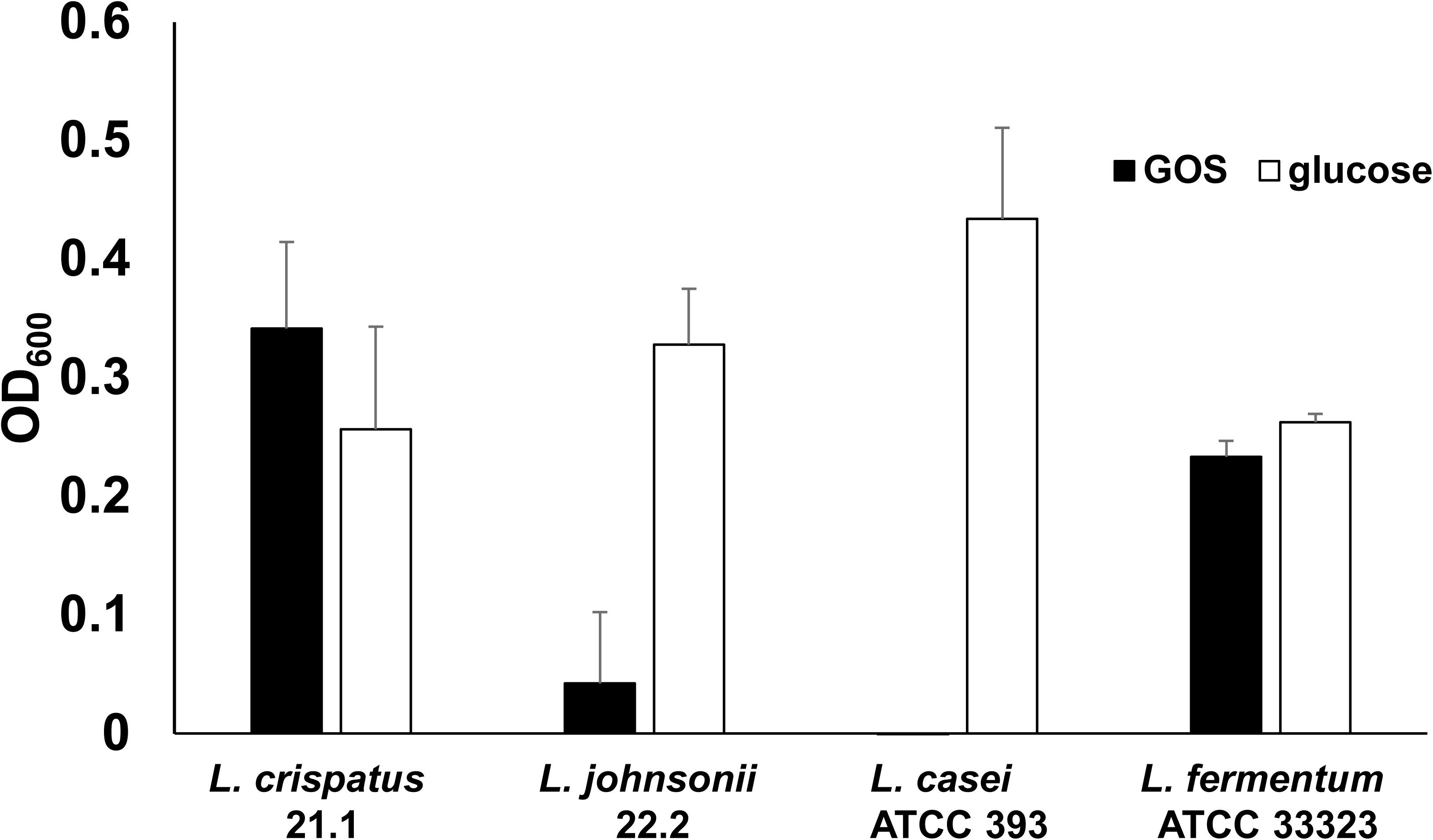
In vitro growth of lactobacilli on galacto-oligosaccharides. Utilization of DP2+ GOS by *L. crispatus* and *L. johnsonii* isolates and *Lactobacillus* type strains. The OD600 recorded after incubation with basal medium was subtracted from that obtained by incubation with basal medium plus DP2+GOS. A positive control with glucose as carbon source was included. The error bars represent ±SD of three biological replicates.

**Figure S3.**
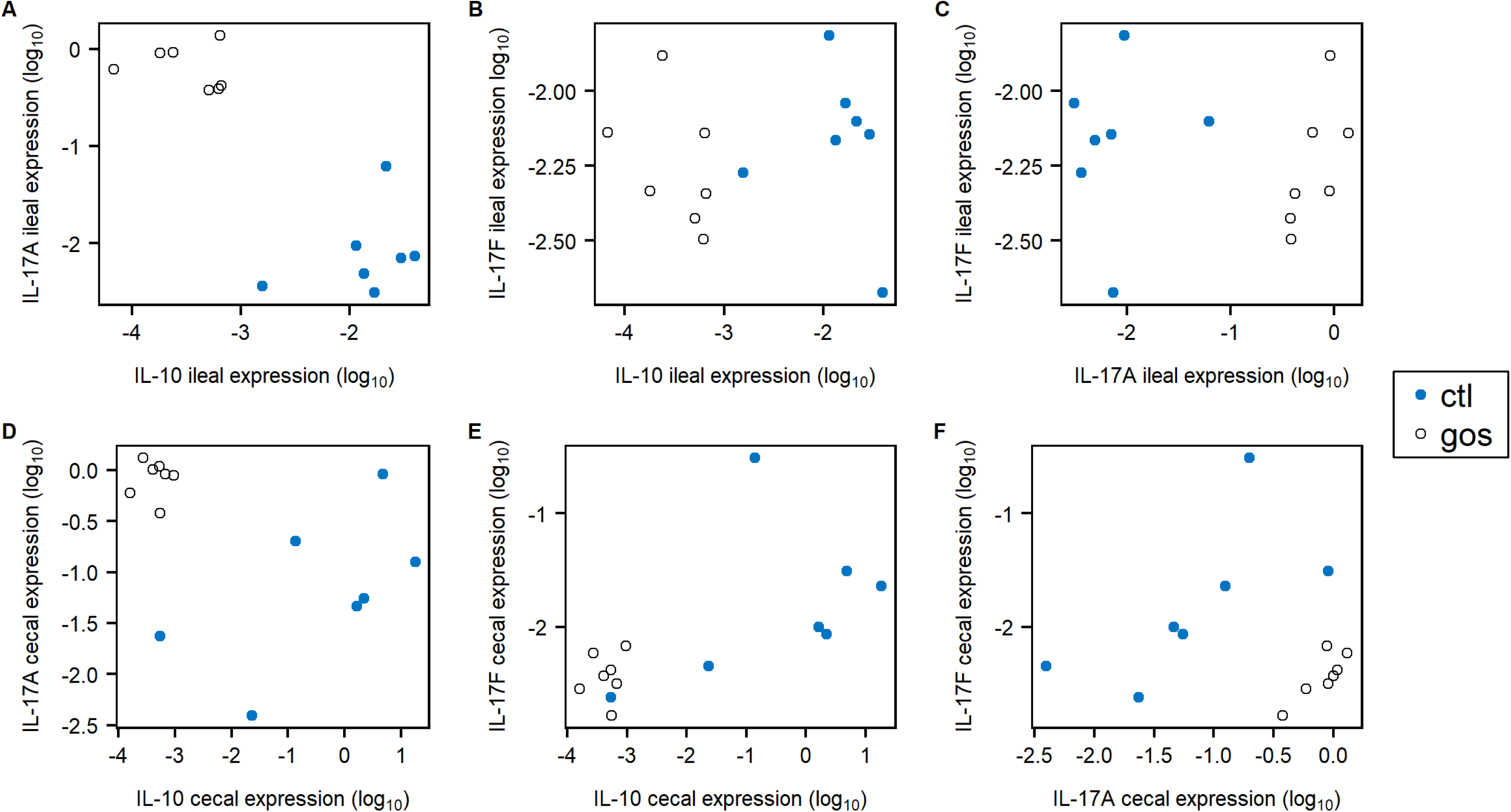
Hematoxylin- and eosin-stained histological examination section of ileal tissues. Tissue sections were made from ileum of 8 da chicken fed a control diet (panel A1), 8 da chicken fed a GOS diet (A2), 35 da chicken fed a control diet (A3), and 35 da chicken fed a GOS diet (A4). Graphic representation of histomorphometric measurements (panel B). The villus length and crypt depth were measured from 10 well-orientated villi for 3 birds from each diet group at each time point. The villus length was measured from the tip of the villus to the opening of the crypt (B1), the crypt depth was measured from the opening to the base of the crypt (B2).

**Figure S4.**
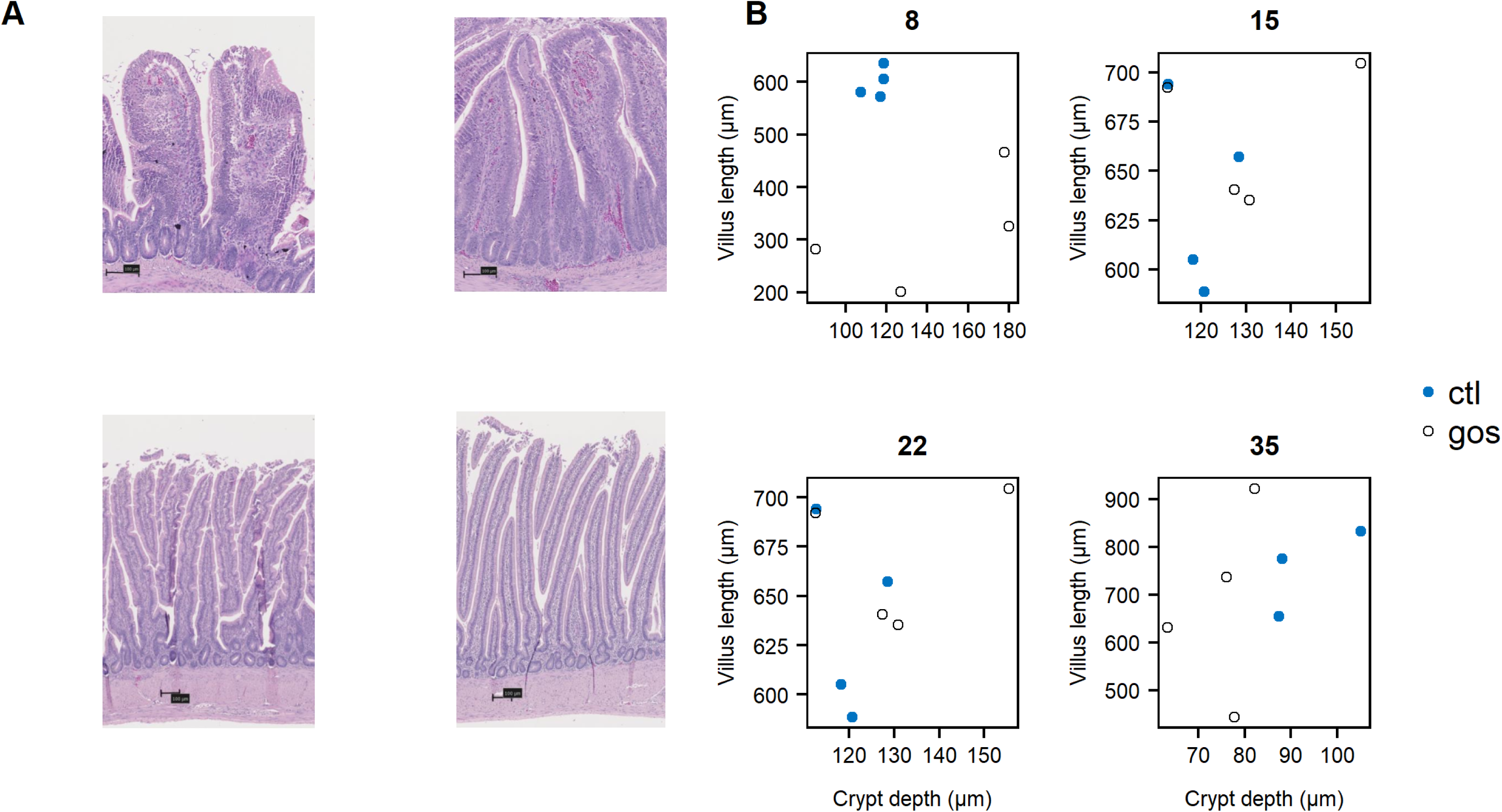
Differential expression in cecal and ileal tissues. Juvenile IL-17A, IL-17F and IL-10 differential gene expression relative to GAPDH at 8 da in ileal (panels A, B, C) and cecal (panels D, E, F) tissues of GOS diet birds (black) compared to control diet birds (blue).

**Figure S5.**
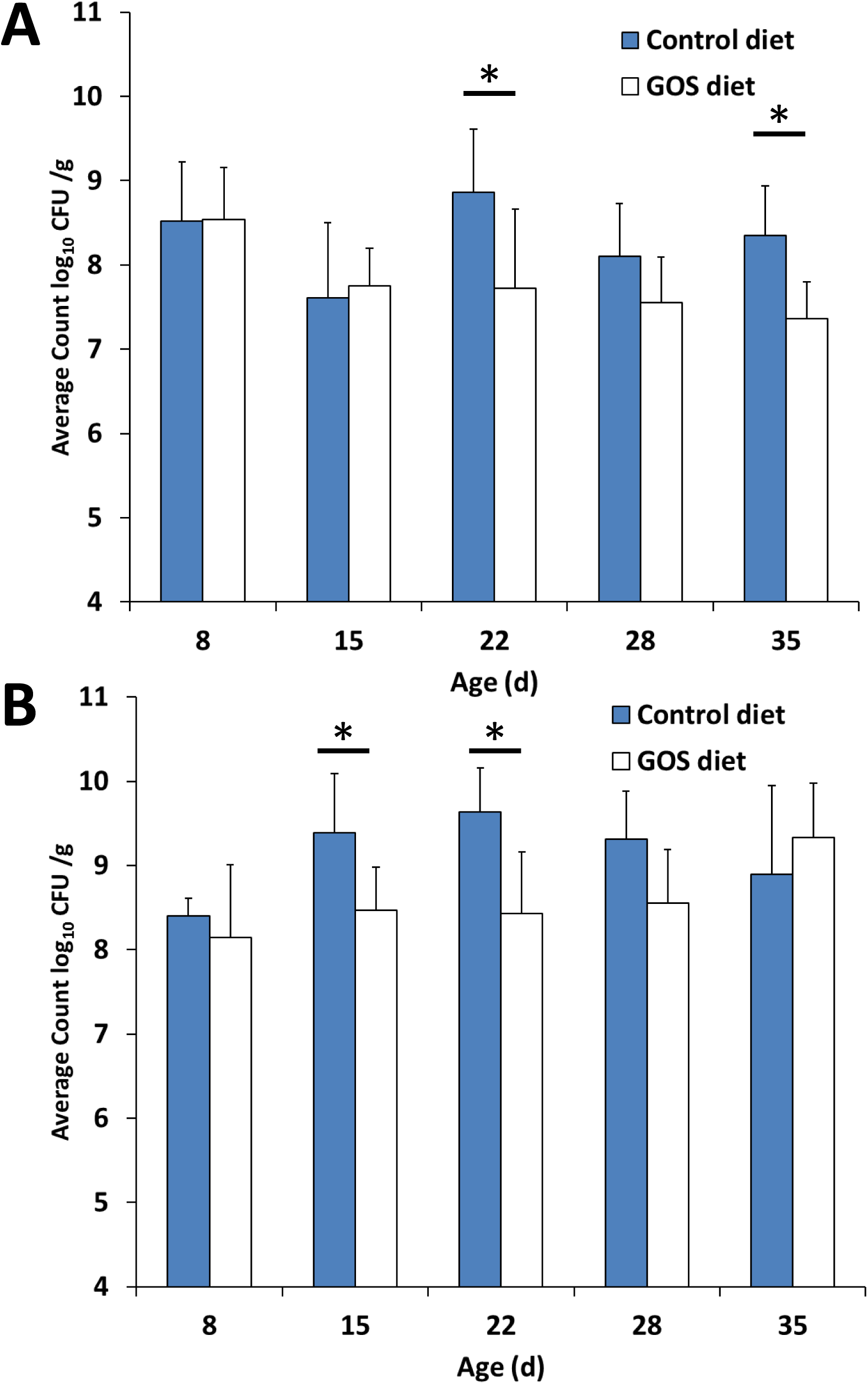
Culture enumeration of Lactic Acid Bacteria and coliforms in cecal contents comparing the two diet regimes. Panels A) Coliform counts B) Lactic Acid Bacteria counts. * indicates p <0.05 (ANOVA).

**Figure S6.**
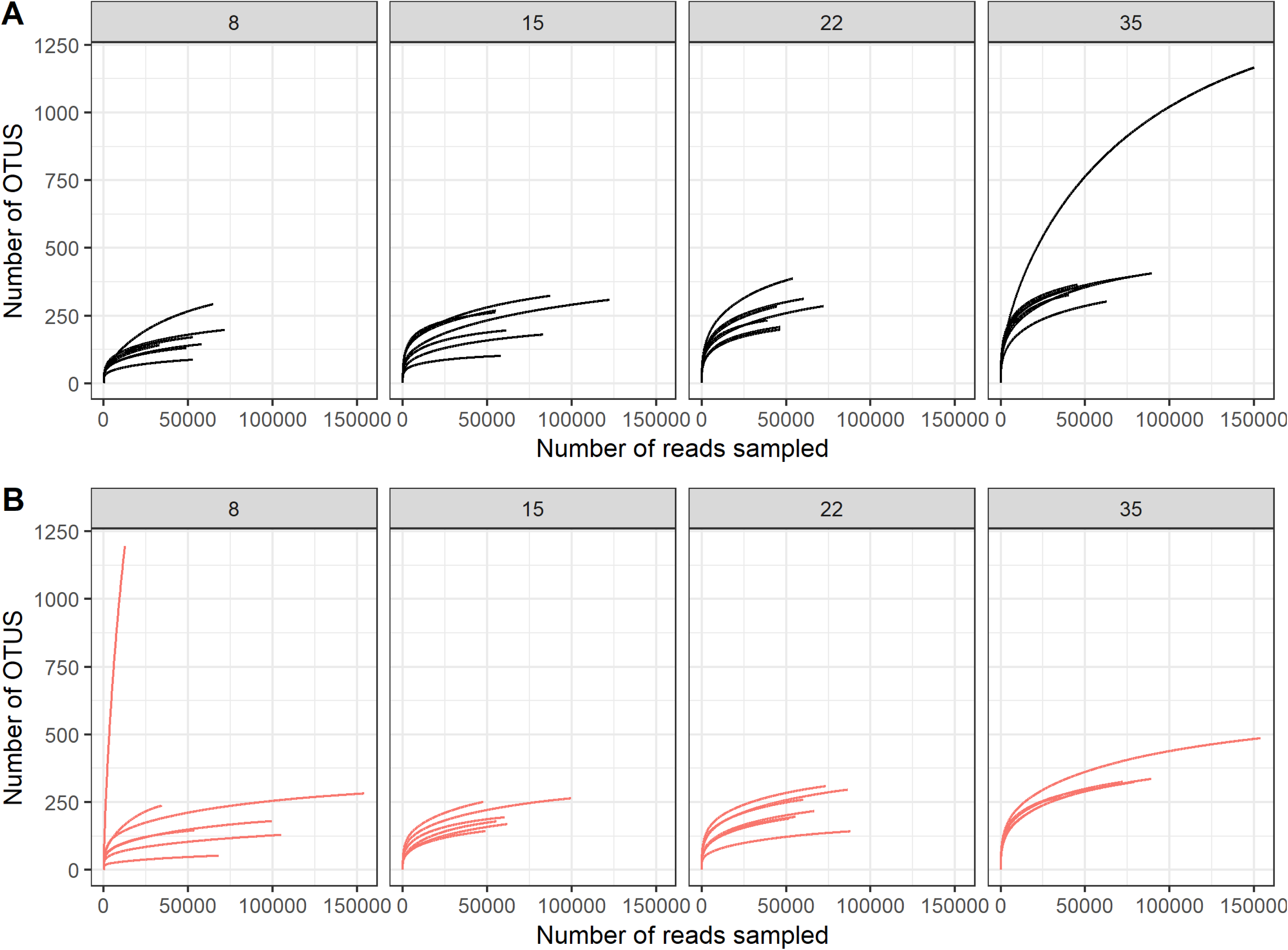
Rarefaction curves indicating coverage of cecal bacterial communities. 16S rDNA bacterial community from panels: A) Control diet 8-35 day of age as indicated above each panel (black); B) GOS-supplemented diet 8-35 day of age as indicated above each panel (red).

**Table S1.** Summary of the functional gene contents of L. johnsonii and L. crispatus isolates related to GOS utilization and host colonization.

| Gene | *L. johnsonii*<br>Locus tag<br>E6A54_ | *L. crispatus*<br>Locus tag<br>E6A57_ | Function | Reference |
| --- | --- | --- | --- | --- |
| *lacS* | 06610* | 07265 | Lactose permease | 39 |
| *lacA* | 06605 | 07260 | β-galactosidase | 39 |
| *lacL/M* | 06620<br>06625 | 07275<br>07280 | β-galactosidase | 34 |
| *lacE/F* | – | 07225 | PTS lactose<br>transporter | 34 |
| *lacG* | – | – | Phospho-<br>β-galactosidase | 34 |
| *mucBP* | 04955<br>08660*<br>09625 | 06095<br>07570<br>08180<br>01350 | Adhesion | 28 |
| *fbpA* | 04640 | 05230 | Adhesion | 28 |
| *cbsA/B* |  | 00840 | S-layer | 32 |
| *apf1/2* | 07620 |  | Aggregation factor | 30 |
| *epsA-E* | 05690-05730 | 08695-08755 | Exopolysaccharide | 29 |
| Bacteriocin-leader |  | 09005 | Bacteriocin | 28 |
| *hly* | 02750 | 00185<br>05290<br>07800<br>10185 | Helveticin J | 28 |
| *bsh* | 00415 | 08515 | Bile salt hydrolase | 31 |

\*internal stop codons present

**Table S2.**
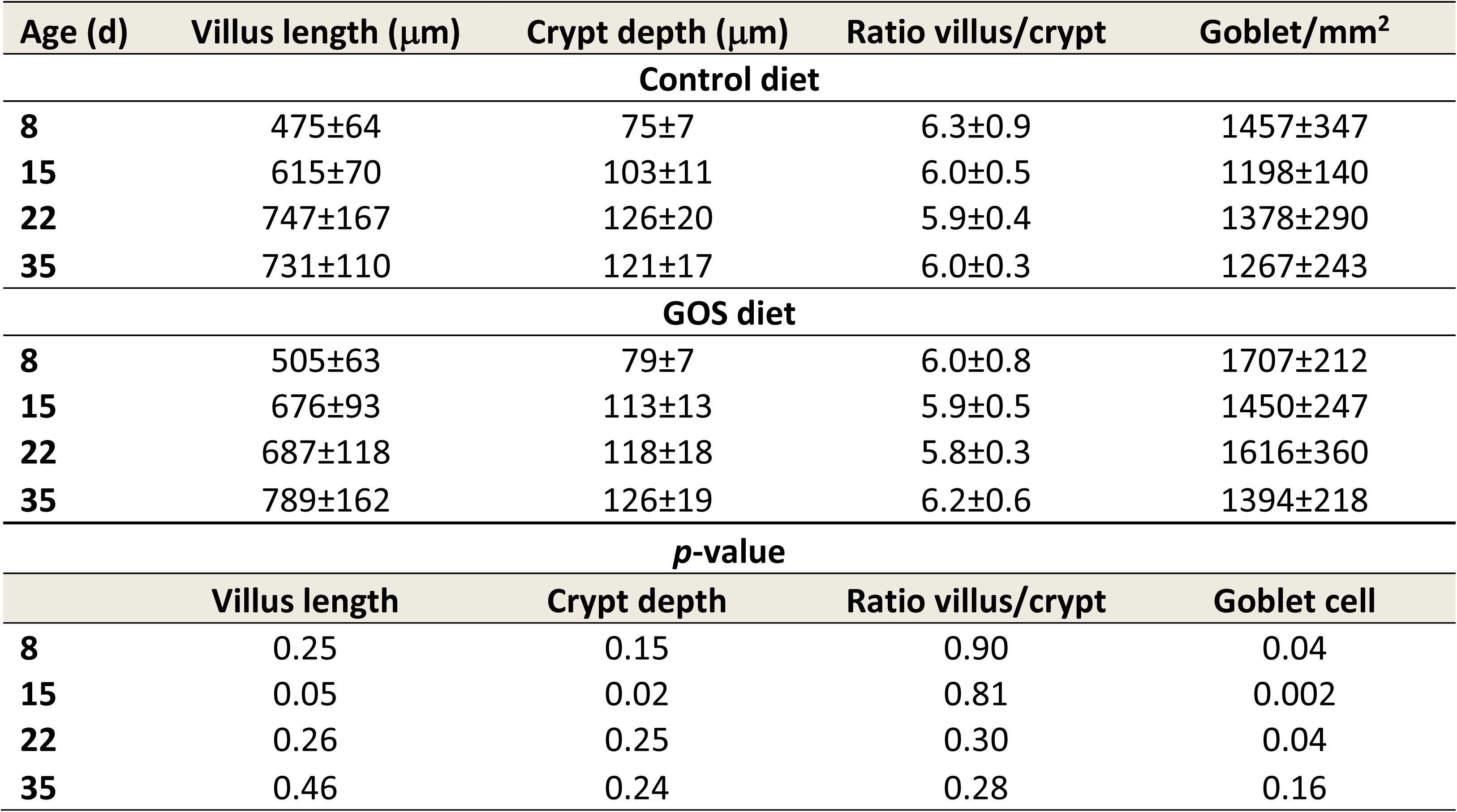
Ileal gut morphometrics: villus length, crypt depth and goblet cell density.

**Table S3.** Modulation of cecal and ileal innate immune responses to dietary GOS in broiler chickens. The table reports cytokine and chemokine gene expression recorded as fold change (FC) for GOS-fed birds relative to the control diet birds at the same age. Corresponding probabilities (*p*) were calculated from ΔCt-values using non-parametric Kruskal-Wallis tests, where differences were considered significant at p<0.05.

**Modulation of cecal and ileal innate immune responses to dietary GOS in broiler chickens.**
|  | **Ceca** |  |  |  |  |  |  |  | **Ileum** |  |  |  |  |  |  |  |
| --- | --- | --- | --- | --- | --- | --- | --- | --- | --- | --- | --- | --- | --- | --- | --- | --- |
|  | **8 da** |  | **15 da** |  | **22 da** |  | **35 da** |  | **8 da** |  | **15 da** |  | **22 da** |  | **35 da** |  |
|  | **FC** | ***p*-value** | **FC** | ***p*-value** | **FC** | ***p*-value** | **FC** | ***p*-value** | **FC** | ***p*-value** | **FC** | ***p*-value** | **FC** | ***p*-value** | **FC** | ***p*-value** |
| **IFNγ** | 0.4 | 0.3 | 11 | 0.05 | 1 | 0.7 | 5 | 0.8 | 0.3 | 0.5 | 1 | 0.7 | 1 | 0.4 | 2 | 0.2 |
| **IL-1β** | 0.1 | 0.9 | 4 | 0.7 | 0.2 | 0.5 | 16 | 0.5 | 1 | 0.5 | 0.1 | 0.2 | 1 | 0.8 | 5 | 0.1 |
| **IL-4** | 112 | 0.5 | 1 | 0.5 | 0.3 | 0.6 | 12 | 0.5 | 0.0001 | 0.3 | 0.02 | 0.02 | 1 | 0.5 | 2 | 0.1 |
| **IL-10** | 0.0001 | 0.01 | 45 | 0.2 | 0.2 | 0.7 | 0.1 | 0.7 | 0.0005 | 0.002 | 3 | 0.7 | 0.3 | 0.3 | 3 | 0.02 |
| **IL-6** | 0.0003 | 0.3 | 1 | 0.03 | 0.1 | 0.4 | 13 | 0.7 | 0.1 | 0.4 | 1 | 0.5 | 1 | 0.9 | 1 | 0.02 |
| **CXCLi-1** | 0.3 | 0.7 | 1 | 0.9 | 0.2 | 0.02 | 2 | 0.9 | 0.3 | 0.7 | 2 | 0.3 | 36 | 0.01 | 0.2 | 0.2 |
| **CXCLi-2** | 0.4 | 0.7 | 7 | 0.2 | 0.2 | 0.03 | 1 | 0.4 | 0.5 | 0.3 | 4 | 0.1 | 36 | 0.02 | 5 | 0.2 |
| **IL-17A** | 5 | 0.01 | 23 | 0.004 | 1 | 0.1 | 52 | 0.002 | 51 | 0.002 | 4196 | 0.003 | 1 | 0.4 | 0.01 | 0.003 |
| **IL-17F** | 0.1 | 0.04 | 0.1 | 1 | 0.1 | 0.4 | 1 | 0.9 | 1 | 0.4 | 19 | 0.004 | 1 | 0.7 | 0.2 | 0.002 |

**Table S4.**
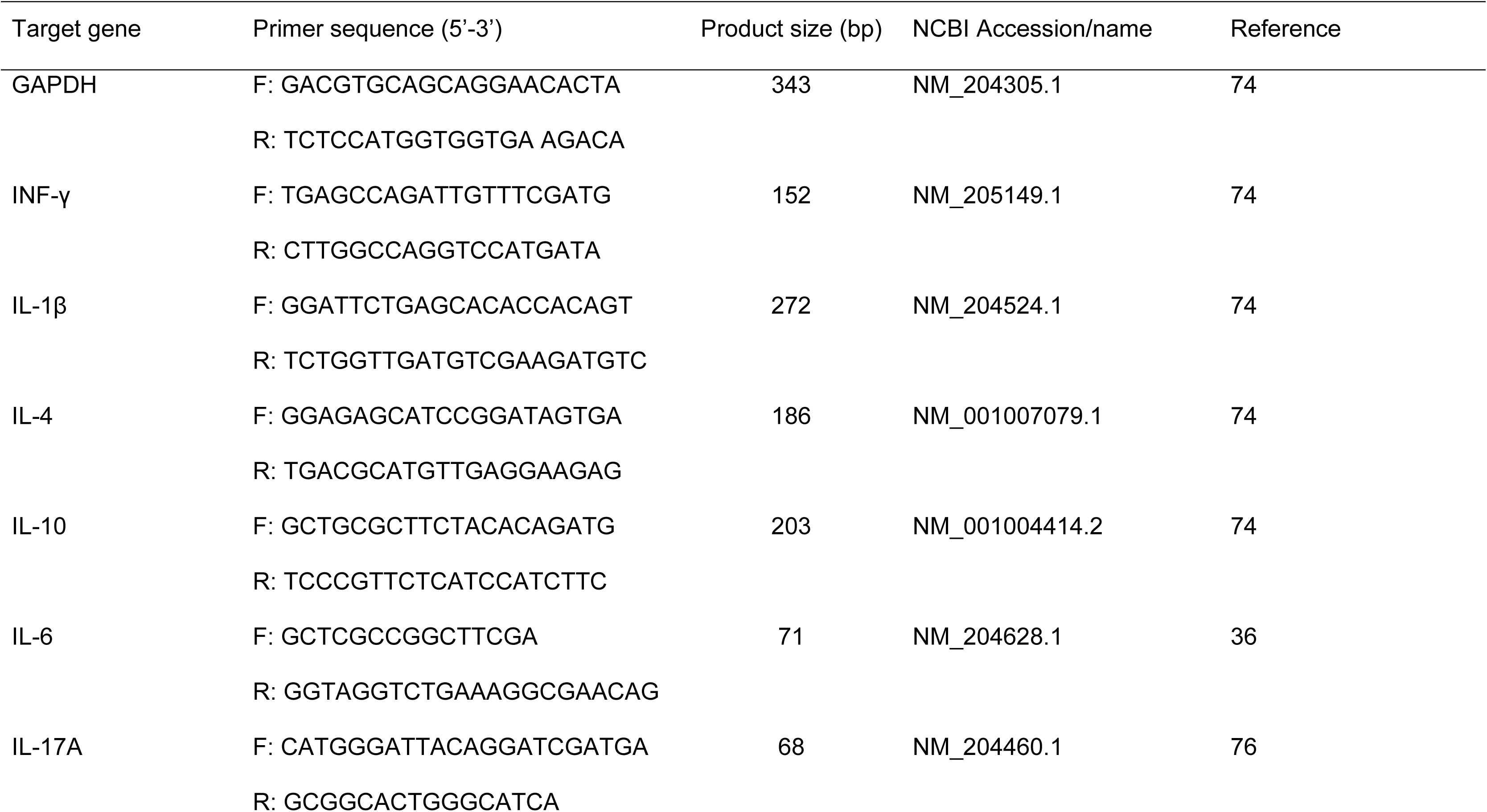
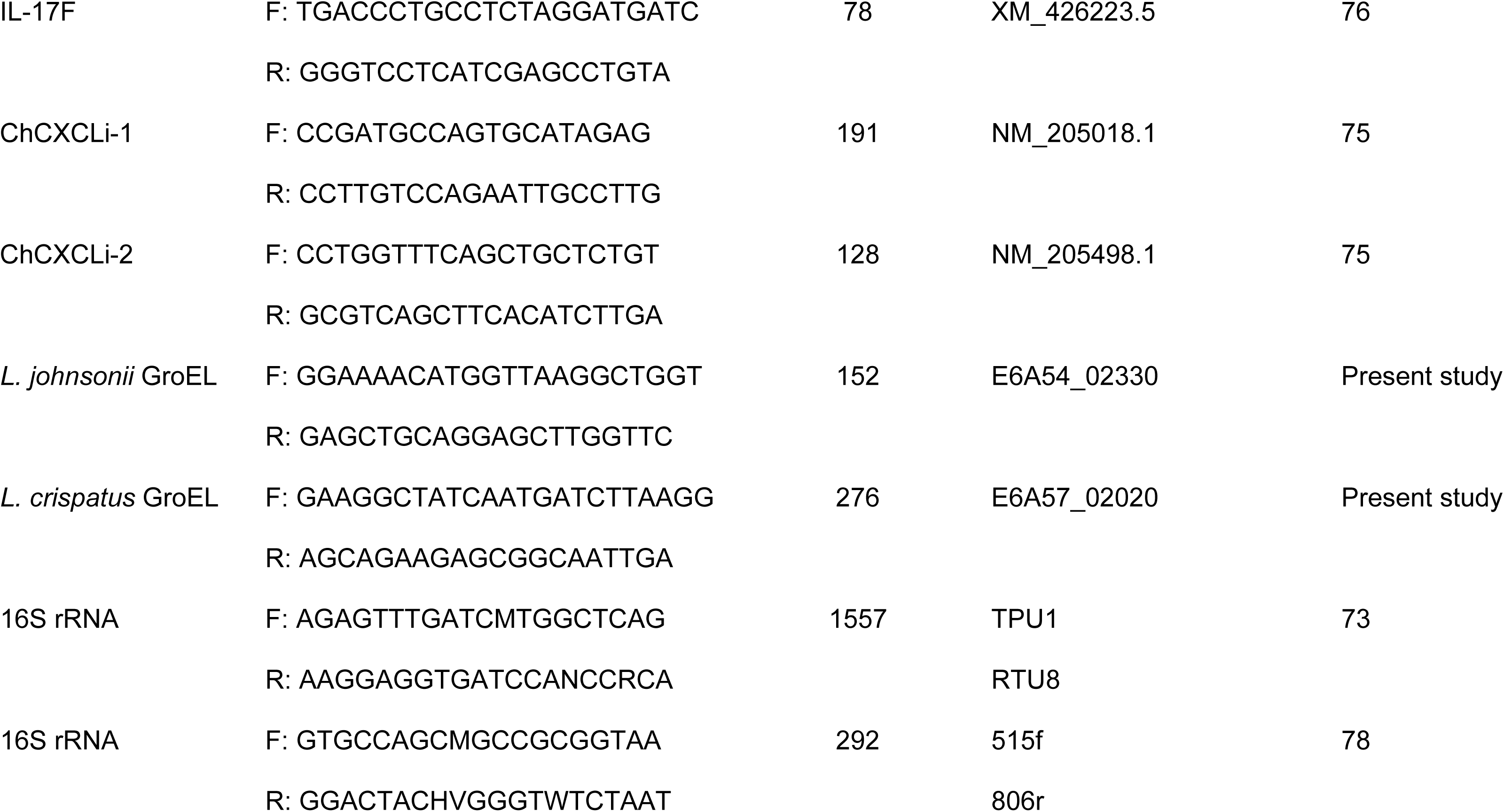
Primer sequences used for PCR. Primer sequences used in this trial with citation.

## REFERENCES

1. Scanes C. 2007. The global importance of poultry. Poult Sci 86:1057–1058,

2. Mottet A, Tempio G. 2017. Global poultry production: Current state and future outlook and challenges. World Poult Sci J 73:245–256.

3. de Vries M, de Boer IJ. 2010. Comparing environmental impacts for livestock products: A review of life cycle assessments. Livest Sci 128:1–11.

4. Zuidhof M, Schneider B, Carney V, Korver D, Robinson F. 2014. Growth, efficiency, and yield of commercial broilers from 1957, 1978, and 2005. Poult Sci 93:2970–2982.

5. Tallentire C, Leinonen I, Kyriazakis I. 2018. Artificial selection for improved energy efficiency is reaching its limits in broiler chickens. Sci Rep 8:1168. DOI:10.1038/s41598-018-19231-2

6. Castanon J. 2007. History of the use of antibiotic as growth promoters in European poultry feeds. Poult Sci 86:2466–2471.

7. Van Immerseel F, Eeckhaut V, Moore RJ, Choct M, Ducatelle R. 2017. Beneficial microbial signals from alternative feed ingredients: A way to improve sustainability of broiler production? Microb Biotech 10:1008–1011.

8. Yadav S, Jha R. 2019. Strategies to modulate the intestinal microbiota and their effects on nutrient utilization, performance, and health of poultry. J Anim Sci Biotech 10:2. https://doi.org/10.1186/s40104-018-0310-9

9. Sethiya NK. 2016. Review on natural growth promoters available for improving gut health of poultry: An alternative to antibiotic growth promoters. Asian J Poult Sci 10:1–29.

10. Thacker PA. 2013. Alternatives to antibiotics as growth promoters for use in swine production: A review. J Anim Sci Biotech 4:35. doi.org/10.1186/2049-1891-4-35

11. Huang P, Zhang Y, Xiao K, Jiang F, Wang H, Tang D, Liu D, Liu B, Liu Y, He X, Liu H, Liu X, Qing Z, Liu C, Huang J, Ren Y, Yun L, Yin L, Lin Q, Zeng C, Su X, Yuan J, Lin L, Hu N, Cao H, Huang S, Guo Y, Fan W, Zeng J. 2018 The chicken gut metagenome and the modulatory effects of plant-derived benzylisoquinoline alkaloids. Microbiome 6:211. doi.org/10.1186/s40168-018-0590-5

12. Erickson KL, Hubbard NE. 2000. Probiotic immunomodulation in health and disease. J Nutr 130:403S–409S.

13. Gadde U, Kim WH, Oh ST, Lillehoj HS. 2017. Alternatives to antibiotics for maximizing growth performance and feed efficiency in poultry: a review. Anim Health Res Rev 18:26–45.

14. Schokker D, Jansman AJ, Veninga G, De Bruin N, Vastenhouw SA, de Bree FM, Bossers A, Rebel JM, Smits MA. 2017. Perturbation of microbiota in one-day old broiler chickens with antibiotic for 24 hours negatively affects intestinal immune development. BMC genomics 18:241. https://doi.org/10.1186/s12864-017-3625-6

15. Stanley D, Hughes RJ, Geier MS, Moore RJ. 2016. Bacteria within the gastrointestinal tract microbiota correlated with improved growth and feed conversion: Challenges presented for the identification of performance enhancing probiotic bacteria. Front Microbiol 7:187. DOI: 10.3389/fmicb.2016.00187

16. Rubio LA. 2019. Possibilities of early life programming in broiler chickens via intestinal microbiota modulation. Poultry Science 98: 695–706.

17. van Leeuwen SS, Kuipers BJH, Dijkhuizen L, Kamerling JP. 2016. Comparative structural characterization of 7 commercial galacto-oligosaccharide (GOS) products. Carbohydrate Research 425:48–58.

18. Van Bueren AL, Mulder M, Van Leeuwen S, Dijkhuizen L. 2017. Prebiotic galactooligosaccharides activate mucin and pectic galactan utilization pathways in the human gut symbiont *Bacteroides thetaiotaomicron*. Sci Rep 7:40478.

19. Jung S, Houde R, Baurhoo B, Zhao X, Lee B. 2008. Effects of galacto-oligosaccharides and a *Bifidobacteria lactis*-based probiotic strain on the growth performance and fecal microflora of broiler chickens. Poult Sci 87:1694–1699.

20. Bednarczyk M, Stadnicka K, Kozłowska I, Abiuso C, Tavaniello S, Dankowiakowska A, Sławińska A, Maiorano G. 2016. Influence of different prebiotics and mode of their administration on broiler chicken performance. Animal 10:1271–1279.

21. Choct M. 2009. Managing gut health through nutrition. Brit Poult Sci 50:9–15.

22. Kraehenbuhl J-P, Neutra MR. 1992. Molecular and cellular basis of immune protection of mucosal surfaces. Physiol Rev 72:853–879.

23. Honda K, Littman DR. 2012. The microbiome in infectious disease and inflammation. Ann Rev Immunol 30:759–795.

24. Atarashi K, Nishimura J, Shima T, Umesaki Y, Yamamoto M, Onoue M, Yagita H, Ishii N, Evans R, Honda K, others. 2008. ATP drives lamina propria T_h_ 17 cell differentiation. Nature 455:808. https://doi.org/10.1038/nature07240

25. Ivanov II, de Llanos Frutos R, Manel N, Yoshinaga K, Rifkin DB, Sartor RB, Finlay BB, Littman DR. 2008. Specific microbiota direct the differentiation of Il-17-producing T-helper cells in the mucosa of the small intestine. Cell Host Microbe 4:337–349.

26. Jian W, Zhu L, Dong X. 2001. New approach to phylogenetic analysis of the genus *Bifidobacterium* based on partial HSP60 gene sequences. Int J Syst Evo Microbiol 51:1633–1638.

27. Junick J, Blaut M. 2012. Quantification of human fecal *Bifidobacterium* species by use of quantitative real-time PCR analysis targeting the *gro*EL gene. Appl Environ Microbiol 78:2613–2622.

28. Ojala T, Kankainen M, Castro J, Cerca N, Edelman S, Westerlund-Wikström B, Paulin L, Holm L, Auvinen P. 2014. Comparative genomics of *Lactobacillus crispatus* suggests novel mechanisms for the competitive exclusion of *Gardnerella vaginalis*. BMC Genomics 5:1070. doi: 10.1186/1471-2164-15-1070.

29. Dertli E, Colquhoun IJ, Gunning AP, Bongaerts RJ, Le Gall G, Bonev BB, Mayer MJ, Narbad A. 2013. Structure and biosynthesis of two exopolysaccharides produced by *Lactobacillus johnsonii* FI9785. J Biol Chem 288:31938–31951.

30. Ventura M, Jankovic I, Walker DC, Pridmore RD, Zink R. 2002. Identification and characterization of novel surface proteins in *Lactobacillus johnsonii* and *Lactobacillus gasseri*. Appl Environ Microbiol 68:6172–6181.

31. O’Flaherty S, Briner Crawley A, Theriot CM, Barrangou R. 2018. The *Lactobacillus* bile salt hydrolase repertoire reveals niche-specific adaptation. mSphere 3: e00140–18. doi: 10.1128/mSphere.00140-18.

32. Sillanpää J, Martínez B, Antikainen J, Toba T, Kalkkinen N, Tankka S, Lounatmaa K, Keränen J, Höök M, Westerlund-Wikström B, Pouwels PH, Korhonen TK. 2000. Characterization of the collagen-binding S-layer protein CbsA of *Lactobacillus crispatus*. J Bacteriol 182:6440–6450.

33. Antikainen J, Anton L, Sillanpää J, Korhonen TK. 2002. Domains in the S-layer protein CbsA of *Lactobacillus crispatus* involved in adherence to collagens, laminin and lipoteichoic acids and in self-assembly. Mol Microbiol 46:381–394.

34. Gänzle MG, Follador R. 2012. Metabolism of oligosaccharides and starch in lactobacilli: a review. Front Microbiol 3:340. doi: 10.3389/fmicb.2012.00340.

35. Wegmann U, Overweg K, Horn N, Goesmann A, Narbad A, Gasson MJ, Shearman C. 2009. Complete genome sequence of *Lactobacillus johnsonii* FI9785, a competitive exclusion agent against pathogens in poultry. J Bacteriol 191:7142–7143

36. Kaiser P, Stäheli P. 2013. Avian Cytokines and Chemokines, pp. 189–204. In Kaspers B, Schat K (ed), Avian Immunology: 2^nd^ ed. Academic Press, Cambridge, Massachusetts.

37. Arsi K, Donoghue AM, Woo-Ming A, Blore PJ, Donoghue DJ. 2015. Intracloacal inoculation, an effective screening method for determining the efficacy of probiotic bacterial isolates against *Campylobacter* colonization in broiler chickens. J Food Prot. 78:209–213.

38. Alizadeh A, Akbari P, Difilippo E, Schols HA, Ulfman LH, Schoterman MH, Garssen J, Fink-Gremmels J, Braber S. 2016. The piglet as a model for studying dietary components in infant diets: effects of galacto-oligosaccharides on intestinal functions. Br J Nutr 115:605–18. doi:10.1017/S0007114515004997.

39. Andersen JM, Barrangou R, Hachem MA, Lahtinen S, Goh YJ, Svensson B, Klaenhammer TR. 2011. Transcriptional and functional analysis of galactooligosaccharide uptake by *lac*S in *Lactobacillus acidophilus*. Proc Natl Acad Sci USA 108:17785–17790.

40. Guinane CM, Kent RM, Norberg S, Hill C, Fitzgerald GF, Stanton C, Ross RP. 2011. Host specific diversity in *Lactobacillus johnsonii* as evidenced by a major chromosomal inversion and phage resistance mechanisms. PLoS One. 6: e18740. doi: 10.1371/journal.pone.0018740.

41. Pridmore, R.D., Berger, B., Desiere, F., Vilanova, D., Barretto, C., Pittet, A.C., Zwahlen, M.C., Rouvet, M., Altermann, E., Barrangou, R., Mollet, B., Mercenier, A., Klaenhammer, T., Arigoni, F., and Schell, M.A. 2004. The genome sequence of the probiotic intestinal bacterium *Lactobacillus johnsonii* NCC 533. Proc Natl Acad Sci USA 101:2512–2517.

42. Altermann E, Russell WM, Azcarate-Peril MA, Barrangou R, Buck BL, McAuliffe O, Souther N, Dobson A, Duong T, Callanan M, Lick S, Hamrick A, Cano R, Klaenhammer TR. 2005. Complete genome sequence of the probiotic lactic acid bacterium *Lactobacillus acidophilus* NCFM. Proc Natl Acad Sci USA 102: 3906–3912

43. Azcarate-Peril MA, Butz N, Cadenas MB, Koci M, Ballou A, Mendoza M, Ali R, Hassan H. 2018. An attenuated *Salmonella enterica* serovar Typhimurium strain and galactooligosaccharides accelerate clearance of *Salmonella* infections in poultry through modifications to the gut microbiome. Appl Environ Microbiol 84:e02526–17. doi: 10.1128/AEM.02526-17.

44. Valladares R, Sankar D, Li N, Williams E, Lai K-K, Abdelgeliel AS, Gonzalez CF, Wasserfall CH, Larkin J III, Schatz D, Atkinson MA, Triplett, EW, Neu J, Lorca GL. 2010. *Lactobacillus johnsonii* N6.2 mitigates the development of type 1 diabetes in BB-DP rats. PLoS One 5:e10507. doi: 10.1371/journal.pone.0010507

45. Marcial GE, Ford AL, Haller MJ, Gezan SA, Harrison NA, Cai D, Meyer JL, Perry DJ, Atkinson MA, Wasserfall CH, Garrett T, Gonzalez CF, Brusko TM, Dahl WJ, Lorca GL. 2017. *Lactobacillus johnsonii* N6.2 modulates the host immune responses: A double-blind, randomized trial in healthy adults. Frontiers in Immunology 8:655. doi: 10.3389/fimmu.2017.00655

46. Fonseca W, Lucey K, Jang S, Fujimura KE, Rasky A, Ting H-A, Petersen J, Johnson CC, Boushey HA, Zoratti E, others. 2017. *Lactobacillus johnsonii* supplementation attenuates respiratory viral infection via metabolic reprogramming and immune cell modulation. Mucosal Immunology 10:1569.

47. DiMarzio M, Rusconi B, Yennawar NH, Eppinger M, Patterson AD, Dudley EG. 2017. Identification of a mouse *Lactobacillus johnsonii* strain with deconjugase activity against the FXR antagonist T-β-MCA. PLoS one 12:e0183564. doi: 10.1371/journal.pone.0183564

48. Wang H, Ni X, Qing X, Zeng D, Luo M, Liu L, Li G, Pan K, Jing B. 2017. Live probiotic *Lactobacillus johnsonii* BS15 promotes growth performance and lowers fat deposition by improving lipid metabolism, intestinal development, and gut microflora in broilers. Frontiers in Microbiology 8:1073. doi: 10.3389/fmicb.2017.01073

49. Qing X, Zeng D, Wang H, Ni X, Liu L, Lai J, Khalique A, Pan K, Jing B. 2017. Preventing subclinical necrotic enteritis through *Lactobacillus johnsonii* BS15 by ameliorating lipid metabolism and intestinal microflora in broiler chickens. AMB Express 7:139. doi: 10.1186/s13568-017-0439-5.

50. Wang H, Ni X, Liu L, Zeng D, Lai J, Qing X, Li G, Pan K, Jing B. 2017. Controlling of growth performance, lipid deposits and fatty acid composition of chicken meat through a probiotic, *Lactobacillus johnsonii* during subclinical *Clostridium perfringens* infection. Lipids Health Dis 16:38. doi: 10.1186/s12944-017-0408-7.

51. Xin J, Zeng D, Wang H, Sun N, Zhao Y, Dan Y, Pan K, Jing B, Ni X. 2019. Probiotic *Lactobacillus johnsonii* BS15 promotes growth performance, intestinal immunity, and gut microbiota in piglets. Probiotics Antimicrob Proteins doi: 10.1007/s12602-018-9511-y.

52. Stanley D, Geier MS, Chen H, Hughes RJ, Moore RJ. 2015. Comparison of fecal and cecal microbiotas reveals qualitative similarities but quantitative differences. BMC Microbiol 15:51. doi: 10.1186/s12866-015-0388-6.

53. La Ragione RM, Narbad A, Gasson MJ, Woodward, M.J. 2004. In vivo characterization of *Lactobacillus johnsonii* FI9785 for use as a defined competitive exclusion agent against bacterial pathogens in poultry. Lett Appl Microbiol 38:197–205.

54. Hütt P, Lapp E, Štšepetova J, Smidt I, Taelma H, Borovkova N, Oopkaup H, Ahelik A, Rööp T, Hoidmets D, Samuel K, Salumets A, Mändar R. 2016. Characterisation of probiotic properties in human vaginal lactobacilli strains. Microb Ecol Health Dis 27:30484. doi: 10.3402/mehd.v27.30484.

55. Kogut MH, Genovese KJ, Swaggerty CL, He H, Broom L 2018 Inflammatory phenotypes in the intestine of poultry: not all inflammation is created equal. Poult Sci 97:2339–2346.

56. Haas R, Smith J, Rocher-Ros V, Nadkarni S, Montero-Melendez T, D’Acquisto F, Bland EJ, Bombardieri M, Pitzalis C, Perretti M, Marelli-Berg FM, Mauro C. 2016. Lactate regulates metabolic and pro-inflammatory circuits in control of T cell migration and effector functions. PLoS Biol 13:e1002202. doi: 10.1371/journal.pbio.1002202.

57. Lee YS, Kim TY, Kim Y, Lee SH, Kim S, Kang SW, Yang JY, Baek IJ, Sung YH, Park YY, Hwang SW, O E, Kim KS, Liu S, Kamada N, Gao N, Kweon MN. 2018. Microbiota-derived lactate accelerates intestinal stem-cell-mediated epithelial development. Cell Host Microbe 24:833–846.e6. doi: 10.1016/j.chom.2018.11.002.

58. Maxwell JR, Zhang Y, Brown WA, Smith CL, Byrne FR, Fiorino M, Stevens E, Bigler J, Davis JA, Rottman JB, Budelsky AL, Symons A, Towne JE. 2015. Differential roles for Interleukin-23 and Interleukin-17 in intestinal immunoregulation. Immunity 43:739–750.

59. Tang C, Kakuta S, Shimizu K, Kadoki M, Kamiya T, Shimazu T, Kubo S, Saijo S, Ishigame H, Nakae S, Iwakura Y. 2018. Suppression of IL-17F, but not of IL-17A, provides protection against colitis by inducing Treg cells through modification of the intestinal microbiota. Nat Immunol 19:755–765.

60. Gaboriau-Routhiau V, Rakotobe S, Lécuyer E, Mulder I, Lan A, Bridonneau C, Rochet V, Pisi A, De Paepe M, Brandi G, Eberl G, Snel J, Kelly D, Cerf-Bensussan N. 2009. The key role of segmented filamentous bacteria in the coordinated maturation of gut helper T cell responses. Immunity 31:677–689.

61. Ivanov II, Atarashi K, Manel N, Brodie EL, Shima T, Karaoz U, Wei D, Goldfarb KC, Santee CA, Lynch SV, Tanoue T, Imaoka A, Itoh K, Takeda K, Umesaki Y, Honda K, Littman DR. 2009. Induction of intestinal Th17 cells by segmented filamentous bacteria. Cell 139:485–498.

62. Schnupf P, Gaboriau-Routhiau V, Gros M, Friedman R, Moya-Nilges M, Nigro G, Cerf-Bensussan N, Sansonetti PJ. 2015. Growth and host interaction of mouse segmented filamentous bacteria *in vitro*. Nature 520:99–103.

63. Schnupf P, Gaboriau-Routhiau V, Sansonetti PJ, Cerf-Bensussan N. 2017. Segmented filamentous bacteria, Th17 inducers and helpers in a hostile world. Curr Opin Microbiol 35:100–109.

64. Tan TG, Sefik E, Geva-Zatorsky N, Kua L, Naskar D, Teng F, Pasman L, Ortiz-Lopez A, Jupp R, Wu HJ, Kasper DL, Benoist C, Mathis D. 2016. Identifying species of symbiont bacteria from the human gut that, alone, can induce intestinal Th17 cells in mice. Proc Natl Acad Sci U S A. 113:E8141–E8150. doi: org/10.1073/pnas.1617460113

65. Granato D, Perotti F, Masserey I, Rouvet M, Golliard M, Servin A, Brassart D. 1999. Cell surface-associated lipoteichoic acid acts as an adhesion factor for attachment of *Lactobacillus johnsonii* La1 to human enterocyte-like Caco-2 cells. Appl Environ Microbiol 65:1071–1077.

66. Granato D, Bergonzelli GE, Pridmore RD, Marvin L, Rouvet M, Corthésy-Theulaz IE. 2004. Cell surface-associated elongation factor Tu mediates the attachment of *Lactobacillus johnsonii* NCC533 (La1) to human intestinal cells and mucins. Infect Immun 72:2160–2169.

67. Bergonzelli GE, Granato D, Pridmore RD, Marvin-Guy LF, Donnicola D, Corthésy-Theulaz IE. 2006. GroEL of *Lactobacillus johnsonii* La1 (NCC 533) is cell surface associated: potential role in interactions with the host and the gastric pathogen *Helicobacter pylori*. Infect Immun 74:425–434.

68. Turpin W, Humblot C, Noordine ML, Thomas M, Guyot JP. 2012. Lactobacillaceae and cell adhesion: genomic and functional screening. PLoS One 7:e38034. doi: 10.1371/journal.pone.0038034.

69. Crhanova M, Hradecka H, Faldynova M, Matulova M, Havlickova H, Sisak F, Rychlik I. 2011. Immune response of chicken gut to natural colonization by gut microflora and to *Salmonella enterica* serovar enteritidis infection. Infect Immun 79:2755–2763.

70. Connerton PL, Richards PJ, Lafontaine GM, O’Kane PM, Ghaffar N, Cummings NJ, Smith DL, Fish NM, Connerton IF. 2018. The effect of the timing of exposure to *Campylobacter jejuni* on the gut microbiome and inflammatory responses of broiler chickens. Microbiome 6:88. doi: 10.1186/s40168-018-0477-5.

71. Johnson TJ, Youmans BP, Noll S, Cardona C, Evans NP, Karnezos TP, Ngunjiri JM, Abundo MC, Lee C-W. 2018. A consistent and predictable commercial broiler chicken bacterial microbiota in antibiotic-free production displays strong correlations with performance. Appl Environ Microbiol 84:e00362–18. doi: 10.1128/AEM.00362-18

72. Zeeshan IU, Sivaloganathan L, McKenna A, Richmond A, Kelly C, Linton M, Stratakos AC, Lavery U, Elmi, A, Wren BW, Dorrell N, Corcionivoschi N, Gundogdu O. 2018. Comprehensive longitudinal microbiome analysis of the chicken cecum reveals a shift from competitive to environmental drivers and a window of opportunity for *Campylobacter*. Frontiers in Microbiology 9:2452 doi: 10.3389/fmicb.2018.02452

73. Ott SJ, Musfeldt M, Ullmann U, Hampe J, Schreiber S. 2004. Quantification of intestinal bacterial populations by real-time PCR with a universal primer set and minor groove binder probes: a global approach to the enteric flora. J Clin Microbiol 42:2566–2572.

74. Nang NT, Lee JS, Song BM, Kang YM, Kim HS, Seo SH. 2011. Induction of inflammatory cytokines and Toll-like receptors in chickens infected with avian H9N2 influenza virus. Vet Res 42:64. doi: 10.1186/1297-9716-42-64.

75. Rasoli M, Yeap SK, Tan SW, Moeini H, Ideris A, Bejo MH, Alitheen NB, Kaiser P, Omar AR. 2014. Alteration in lymphocyte responses, cytokine and chemokine profiles in chickens infected with genotype VII and VIII velogenic Newcastle disease virus. Comp Immunol Microbiol Infect Dis 37:11–21.

76. Reid WD, Close AJ, Humphrey S, Chaloner G, Lacharme-Lora L, Rothwell L, Kaiser P, Williams NJ, Humphrey TJ, Wigley P, Rushton SP. 2016. Cytokine responses in birds challenged with the human food-borne pathogen *Campylobacter jejuni* implies a Th17 response. R Soc Open Sci 3:150541. doi: 10.1098/rsos.150541

77. Livak K.J. and Schmittgen T.D. 2001. Analysis of relative gene expression data using real-time quantitative PCR and the 2-ΔΔCT Method. Methods 24:402–408.

78. Caporaso JG, Lauber CL, Walters WA, Berg-Lyons D, Lozupone CA, Turnbaugh PJ, Fierer N, Knight R. 2011. Global patterns of 16S rRNA diversity at a depth of millions of sequences per sample. Proc Natl Acad Sci U S A 108: Suppl 1:4516–22. doi: 10.1073/pnas.1000080107.

79. Schloss PD, Westcott SL, Ryabin T, Hall JR, Hartmann M, Hollister EB, Lesniewski RA, Oakley BB, Parks DH, Robinson CJ, Sahl JW, Stres B, Thallinger GG, Van Horn DJ, Weber CF. 2009. Introducing mothur: open-source, platform-independent, community-supported software for describing and comparing microbial communities. Appl Environ Microbiol 75:7537–7541

80. Westcott SL, Schloss PD. 2017. OptiClust, an improved method for assigning amplicon-based sequence data to operational taxonomic units. mSphere 2:e00073–17. doi: 10.1128/mSphereDirect.00073-17.

81. Kozich JJ, Westcott SL, Baxter NT, Highlander SK, Schloss PD. 2013. Development of a dual-index sequencing strategy and curation pipeline for analyzing amplicon sequence data on the MiSeq Illumina sequencing platform. Appl Environ Microbiol 79:5112–5120.

82. Segata N, Izard J, Waldron L, Gevers D, Miropolsky L, Garrett WS, Huttenhower C. 2011. Metagenomic biomarker discovery and explanation. Genome Biol 12:R60. doi: 10.1186/gb-2011-12-6-r60.

83. R Core Team. 2019. R: A language and environment for statistical computing. R Foundation for Statistical Computing, Vienna, Austria.

84. Warnes GR, Bolker B, Bonebakker L, Gentleman R, Huber W, Liaw A, Lumley T, Maechler M, Magnusson A, Moeller S, Schwartz M, Venables B. 2016 gplots: Various R Programming Tools for Plotting Data. R package version 3.0.1. https://CRAN.R-project.org/package=gplots.

85. RStudio Team. 2015. RStudio: Integrated development environment for r. RStudio, Inc., Boston, MA.

